# Large-scale exploration of whole-brain structural connectivity in anorexia nervosa: alterations in the connectivity of frontal and subcortical networks

**DOI:** 10.1101/2021.10.05.463197

**Authors:** E. Caitlin Lloyd, Karin E. Foerde, Alexandra F. Muratore, Natalie Aw, David Semanek, Joanna E. Steinglass, Jonathan Posner

**Affiliations:** Department of Psychiatry, Columbia University Irving Medical Center, New York, NY, USA; New York State Psychiatric Institute, New York, New York, NY, USA; Department of Psychiatry, Duke University, Durham, NC, USA

**Author notes:** denotes corresponding author, Corresponding author: Dr Caitlin Lloyd, Address: Department of Psychiatry, Columbia University Irving Medical Center, New York, NY, USA. denotes shared last author (Joanna E. Steinglass, Jonathan Posner).

**Keywords:** anorexia nervosa, connectomics, diffusion magnetic resonance imaging, graph theory, structural connectivity, tractography

## Abstract

**Background:** Anorexia nervosa (AN) is characterized by disturbances in cognition and behavior surrounding eating and weight. The severity of AN combined with the absence of localized brain abnormalities suggests distributed, systemic underpinnings that may be identified using diffusion-weighted MRI (dMRI) and tractography to reconstruct white matter pathways.

**Methods:** dMRI data acquired from female patients with AN (n = 147) and female healthy controls (HC; n = 119), aged 12-40 years, were combined across five studies. Probabilistic tractography was completed, and full cortex connectomes describing streamline counts between 84 brain regions generated and harmonized. Graph theory methods were used to describe alterations in network organization in AN. The network-based statistic tested between-group differences in brain subnetwork connectivity. The metrics strength and efficiency indexed the connectivity of brain regions (network nodes), and were compared between groups using multiple linear regression.

**Results:** Individuals with AN, relative to HC, had reduced connectivity in a network comprising subcortical regions and greater connectivity between frontal cortical regions (p < 0.05, FWE corrected). Node-based analyses indicated reduced connectivity of the left hippocampus in patients relative to HC (p < 0.05, permutation corrected). Severity of illness, assessed by BMI, was associated with subcortical connectivity (p < 0.05, uncorrected).

**Conclusions:** Analyses identified reduced structural connectivity of subcortical networks and regions, and stronger cortical network connectivity, amongst individuals with AN relative to HC. These findings are consistent with alterations in feeding, emotion and executive control circuits in AN, and may direct hypothesis-driven research into mechanisms of persistent restrictive eating behavior.

## Introduction

Anorexia nervosa (AN) is a serious illness characterized by extreme food restriction and disturbances in cognition surrounding eating and weight (1). The illness is associated with high rates of mortality (2), medical and psychiatric morbidity (3, 4), and severe decrements in quality of life (5). Treatment for AN is challenging, with disappointing remission rates and high rates of relapse (6). The severity of the psychiatric symptoms in AN makes it somewhat puzzling that mechanisms of illness have not been more easily localized, despite increasing research into the underlying neural pathology (7). Inconclusive findings from hypothesis-driven research leads to the data-driven approach in this study. We examine brain-wide structural connectivity by leveraging advances in network-based models of the brain, and a state-of-the-art probabilistic approach to diffusion-weighted MRI (dMRI) tractography.

White matter connects spatially distinct brain regions, creating the structural connections that support coordinated neural activation. Thus, white matter connections are the structural architecture underlying neural networks (8, 9), and disturbances in this architecture, or connectivity, may underlie abnormal functioning of neural networks. White matter tracts of the brain can be delineated in vivo using dMRI (10), meaning dMRI can facilitate circuit-based inquiry into disease mechanisms in AN.

To date, several dMRI studies of AN have been conducted and generated interesting findings (for review, see (11)). Despite the promise of this work, studies have been limited in several ways. First, most dMRI studies of AN have focused on regional white matter microstructure. Microstructural measurements provide information about the integrity of white matter within a given brain region, but do not assess region-to-region connectivity. Conversely, diffusion tractography is a dMRI approach that captures connectivity by reconstructing white matter pathways throughout the brain. Several diffusion tractography studies of AN have been reported, but there have been important limitations. Most notably, studies have been constrained by small sample sizes, and consistent findings across studies have not emerged (11-13). The single tractography study that had a relatively large sample (n= 96 AN and 96 HC (14)) used deterministic tractography, an approach that assumes that all white matter fibers within a voxel are oriented in the same direction. White matter often contains complex fiber architectures such as bending or crossing fibers (15, 16). More biologically accurate representations of white matter pathways are provided by probabilistic algorithms able to model multiple fiber orientations per voxel (17). In sum, while dMRI offers a promising tool for studying brain architecture, prior dMRI studies of AN have been limited by methodological approaches.

The current study addresses the aforementioned limitations, and differs from the majority of prior dMRI research by adopting a data driven, or hypothesis-generating, approach. Discovery-based, or data driven, approaches are particularly useful when prior knowledge, upon which hypotheses are formulated, is limited—as is the case with the knowledge of white matter architecture in AN. Whilst testing directed hypotheses can be well suited for identifying disease mechanisms, the narrow lens inherent to hypothesis testing can miss unanticipated but meaningful differences in brain architecture. Data-driven approaches can direct the development of questions concerning the possible underpinnings of disease, which can be subjected to hypothesis testing in subsequent studies.

Leveraging a large sample and techniques to handle multiple comparisons, the current study obviates a central concern of data-driven approaches—generating false-positive results. In particular, a graph theory approach is used to explore structural connectivity in AN. The application of mathematical graph theory in neuroscience has led to a network model of the brain: the structural connectome, which comprises a set of “nodes” (brain regions) and “edges” (pairwise structural connections between all regions of the brain (18)). The approach allows for studying the neural architecture across multiple connections, rather than the strength of a discrete pairwise connection, and has been shown to be highly reliable (19-21). Psychopathology has increasingly been linked to abnormalities at the level of diffuse brain circuits (22), making graph theory approaches particularly compelling. Graph theory tools include network-based and node-based analyses. Network-based methods can identify subnetworks associated with a particular trait, and node-based analyses identify brain regions showing variation in whole-brain connectivity across individuals or groups. These methods provide different information about brain organization, and using them together can provide detailed information about localized abnormalities in AN that may be probed further. Graph theory approaches have been applied in AN (14, 23-25) however this study is the first to simultaneously probe the connectivity of subnetworks and nodes.

## Methods

Analysis of dMRI data collected from 266 participants via five cross-sectional studies (12, 26-29) was undertaken. Data from a subsample of participants from one study (n= 21 AN and 18 HC) were included in a prior publication (12).

### Participants

Participants were 147 females who met DSM-5 criteria for AN, and 119 age-matched female HC. Participants were between the ages of 12 and 40 years. Exclusion criteria were the same across studies: current substance use disorder, lifetime psychotic disorder, history of a neurological disorder/injury, estimated IQ less than 80, and contraindications to MRI. Additional exclusion criteria for HC included any current/past major psychiatric illness, serious or unstable medical condition, and body mass index (BMI) outside of the normal range (18.5-25 kg/m^2^). AN patients were free from psychotropic medication use at the time of MRI scanning, except for 11 patients taking antidepressant medication. Eating disorder diagnoses were made by the Eating Disorder Examination (30) or Eating Disorder Diagnostic Assessment for DSM-5 (31). Co-occurring diagnoses were assessed by Structured Clinical Interview for DSM-IV or DSM-5 for adults (32, 33), and by the Schedule for Affective Disorders and Schizophrenia for School-Aged Children for adolescents (34). Height and weight were measured on the day of the MRI scan by stadiometer and balance beam scale respectively. Each study was approved by the New York State Psychiatric Institute (NYSPI) Institutional Review Board. Researchers provided a complete description of the study to participants who provided written informed consent (adults) or written assent with parental consent (adolescents).

Amongst the patients, 84 (57.1%) were diagnosed with AN restricting subtype. Adults with AN were receiving specialized inpatient eating disorder treatment at NYSPI. Adolescents were receiving inpatient or outpatient treatment. With a few exceptions, scanning occurred shortly after patients’ diagnostic assessment (median delay = 2.44 days, interquartile range = 0.91 to 7.94 days).

### MRI procedures

#### Scanning parameters

The MRI scanner and DWI scanning parameters are described in the Supplement (**Table S1**) for each of the five studies. All DWI acquisitions were single shell.

### MRI Image processing

#### Anatomical images

Structural images were processed using the automated Freesurfer (version 6.0.0) pipeline (35, 36). Grey matter regions were parsed into 84 distinct regions based on the Desikan-Killiany atlas (37), given structural connectivity metrics derived from this parcellation have established reliability (19).

The parcellated image was used to generate a five-tissue type image, in which white matter, cortical and subcortical grey matter, cerebral spinal fluid (CSF), and pathological tissue are distinguished. Anatomical images were warped into diffusion space, and resampled to a voxel size of 1.25 mm^3^.

#### Diffusion images

Each scan was visually checked for artifacts prior to image pre-processing, and volumes containing artifacts removed. Participants’ data were excluded if more than 25% of the scan volumes were removed (n = 3 AN and 0 HC). A fully automated image pre-processing pipeline implemented in MRtrix3 (38) was applied to the data, including: denoising, gibbs-artifact removal, B0 susceptibility-induced distortion correction, eddy current and motion correction, and B1 field inhomogeneity correction. Preprocessed dMRI data were resampled to an isotropic voxel size of 1.25 mm^3^. Motion was quantified with average framewise displacement across the dMRI scan and did not differ between participant groups: HC=0.16 (SD=0.1), AN = 0.18 (SD=0.15); t(2) = 1.09, p = 0.273. Average framewise displacement by diagnostic group and study is reported in the Supplement (**Table S2**).

### Reconstruction of White Matter - Probabilistic Tractography

To delineate the white matter pathways of the brain, anatomically-constrained probabilistic tractography was completed. A standard MRtrix3 pipeline (38), described in the Supplement, reconstructed plausible white matter pathways by propagating along estimates of voxel fiber orientation distributions. Voxel fiber distributions were recovered using multi-shell multi-tissue constrained spherical deconvolution (39), a method that minimizes bias due to partial volume effects (resulting from grey matter loss in AN). The reconstruction algorithm generated 20,000,000 streamlines between 10 and 250 mm. These streamlines were filtered (retaining 5,000,000 of them) using the SIFT algorithm, which maximizes the fit between the reconstructed streamlines and dMRI data.

### Constructing structural connectomes

To create brain networks (i.e., structural connectomes), a node was assigned to each of the 84 regions from the anatomical parcellation. For each participant, the number of streamlines between all possible pairs of regions, or nodes (Ni, Nj), was counted to provide edge values in individual brain networks. Streamline endpoints were mapped to nodes within a 4mm radius of the endpoint.

To remove effects of differences in scanner/scan parameters across and within studies, individual brain networks were harmonized using the R implementation of neuroCombat (40, 41).

Diagnosis, age and BMI were specified as covariates in the harmonization model to retain effects of biologically relevant variables in the brain connectivity data whilst removing variance introduced by scan differences. In exploratory analyses probing subtype differences in structural connectivity, the harmonization model was applied to data from AN patients only, including subtype as an additional covariate.

Following recent recommendations, brain networks were not thresholded prior to completing analyses (42).

### Data analysis

Streamline counts between brain regions constituted the measure of connectivity in all analyses. Network-based statistic (43)

Subnetwork connectivity within the structural connectome was compared between AN and HC groups using the network-based statistic method implemented in Matlab (https://sites.google.com/site/bctnet/comparison/nbs; (44)). This method assesses whether groups show altered streamline counts in constellations of connected tracts, or subnetworks, and is fully described in the Supplement. The analysis was completed for the contrasts HC > AN, and AN > HC; p-values were Bonferroni-corrected for two tests. Models were adjusted for age, motion and study. Study was included as a covariate given harmonization may not fully remove batch effects.

#### Node-based analyses

*Strength* and *efficiency*, which describe connectivity properties of brain regions (nodes), were computed for each network node (see the Supplement for further details), using the Matlab Brain Connectivity Toolbox (44, 45). These are standard metrics in studies of brain network structure (45) that provide complementary information about brain network organization. Strength provides an indicator of node centrality, or the extent to which a node influences and is influenced by other network nodes. Efficiency describes the ease with which information can move from one region to another within a network, “ease” referring to fewer intermediary nodes (46). *Global efficiency*, the ease with which information can move from one region to any other on average across the whole brain (46) was also calculated, to examine widespread alterations in brain structure in AN.

Node characteristics were compared between individuals with AN and HC in linear regression models adjusted for age, motion and study. Permutation tests established statistical significance and corrected for multiple comparisons across the 168 tests (84 nodes, two node metrics) using the minP method and the R package Permuco (47). 10,000 permutations for each test were specified; the Freedman-Lane procedure managed effects of nuisance variables (age, motion and study). A linear regression model adjusted for age, motion and study compared AN and HC groups on the global efficiency metric.

The tractography, connectome generation, node metric calculation and analysis procedures are detailed in **Figure 1**.

**Figure 1:**
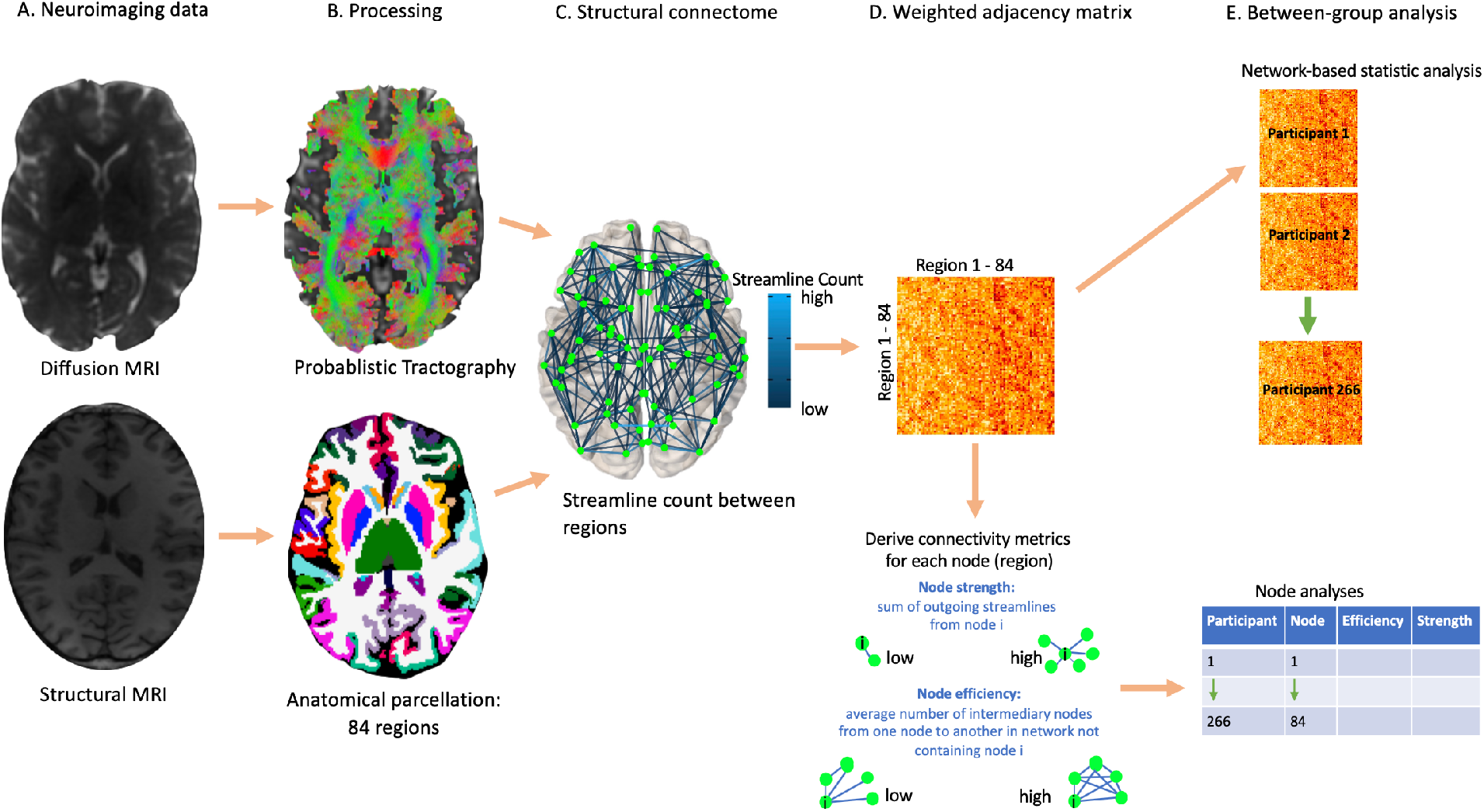
Neuroimaging and analysis methods For each participant, a diffusion MRI and structural MRI scan were acquired (panel A). White matter tracts in the brain were reconstructed using a probabilistic tracking algorithm applied to diffusion MRI data, and the structural brain image was parcellated into 84 regions (panel B). Combining the tractography output with the brain parcellation allowed the creation of a structural connectome for each participant, which described the number of streamlines between each pair of regions (panel C). This information was represented in a weighted adjacency matrix, which was used to calculate node metrics (panel D). Anorexia nervosa and healthy control groups were compared on the strength of structural connections and node metrics, across the whole brain (panel E).

The main analyses probing group differences in brain network attributes were repeated using unharmonized data, to verify whether findings were consistent across different methods for combining data from the five studies.

### Exploratory analyses

Subnetwork components that significantly differed between groups were tested for their association with illness severity, indexed by BMI on the day of the scan (rather than at diagnostic assessment) and illness duration, within the AN group. For each component reaching statistical significance, the mean number of streamlines across edges of these components was calculated. To combine BMI measurements across adults (aged over 19 years) and adolescents (aged 19 years and below), standardized scores (mean centred and scaled by the standard deviation) were calculated in the two groups, using BMI for adults, and BMI percentile for adolescents. These standardized scores were combined to create the standardized BMI variable. Network component connectivity metrics were regressed onto standardized BMI and illness duration, in separate models. Using the same method, node metrics that differed between groups were tested for their association with clinical variables amongst individuals with AN. All statistical models were adjusted for motion, age and study. Models involving the regression of illness duration were additionally adjusted for standardized BMI. Model predictors and outcomes (aside from the already standardized BMI variable) were mean centred and scaled.

Component connectivity and node metrics that differed between AN and HC groups were compared 1) between AN subtypes, in linear regression models adjusted for motion, age, study, and standardized BMI, and 2) between AN and HC in analyses including a) only adolescents, and b) only adults (in linear regression models adjusted for motion, age, and study).

Exploratory analyses were not corrected for multiple comparisons.

### Code and data availability

Code for dMRI data preprocessing, tractography, and analysis steps is available on github: https://github.com/CaitlinLloyd/Anorexia.Nervosa_Tractography. Data for between-group differences in streamline count between all pairs of regions are available (Supplementary Tables).

## Results

### Participant characteristics

Participant characteristics are displayed in **Table 1**. The Supplement (**Table S2**) contains a summary of results by study and diagnosis. Groups were well-matched in age, and differed as expected in BMI at diagnostic assessment and all clinical variables.

**Table 1:** Demographic and clinical characteristics

| Variable | HC<br>(n = 119) | AN<br>(n = 147) |
| --- | --- | --- |
| | Mean ± SD<br>(range) | Mean ± SD<br>(range) |
| Age (years) | 21.03 ± 5.11<br>(12, 39) | 20.34 ± 6.21<br>(12, 40) |
| BMI (kg/m<sup>2</sup>) scan day* | 21.49 ± 1.69<br>(18.7, 25.0)<br>(n = 65) | 16.1 ± 1.98<br>(11.20, 19.90)<br>(n = 60) |
| BMI percentile (%) at scan day* | 47.74 ± 23.38<br>(6.1, 87.4)<br>(n = 54) | 9.60 ± 10.5<br>(0, 45.6)<br>(n = 87) |
| BMI (kg/m<sup>2</sup>) at diagnostic assessment* | NA | 15.92 ± 1.78<br>(11.72, 18.73)<br>(n = 58)<sup>c</sup> |
| BMI percentile (%) at diagnostic assessment* | NA | 8.06 ± 9.32<br>(0, 39.6)<br>(n = 86)<sup>c</sup> |
| EDEQ Global score* | 0.27 ± 0.39<br>(0, 2.17) | 3.9 ± 1.54<br>(0, 6) |
| Illness Duration (months) | NA | 58.15 ± 69.61<br>(0, 324) |
|  | N |  |
| Comorbid anxiety, obsessive-compulsive, or major depressive disorder | NA | n=59 (40.1 %) |
| <sup>a</sup> BMI information presented for participants **over 19** |  |  |
| <sup>b</sup> BMI percentile information presented for participants **19 and under** |  |  |
| <sup>c</sup> Data for BMI at diagnostic assessment is missing for 2 adults with AN, and 1 adolescent with AN |  |  |
| * p-value for between group difference < 0.001 |  |  |
| BMI: Body mass index; EDEQ: Eating disorder examination questionnaire |  |  |

### Network-based statistic

The network-based statistic indicated one component with stronger connectivity amongst HC compared to AN (**Figure 2, Panel A**). This component included 10 connections between subcortical regions (p = 0.041). Conversely, the network-based statistic indicated one component that was stronger amongst individuals with AN relative to HC (**Figure 2, Panel B**). This component included 8 connections between regions generally located in the frontal cortex (p = 0.043). Details of average connectivity within the subcortical and cortical components, by study and diagnosis, are presented in the Supplement (**Table S3, Figures S1 and S2**).

**Figure 2.**
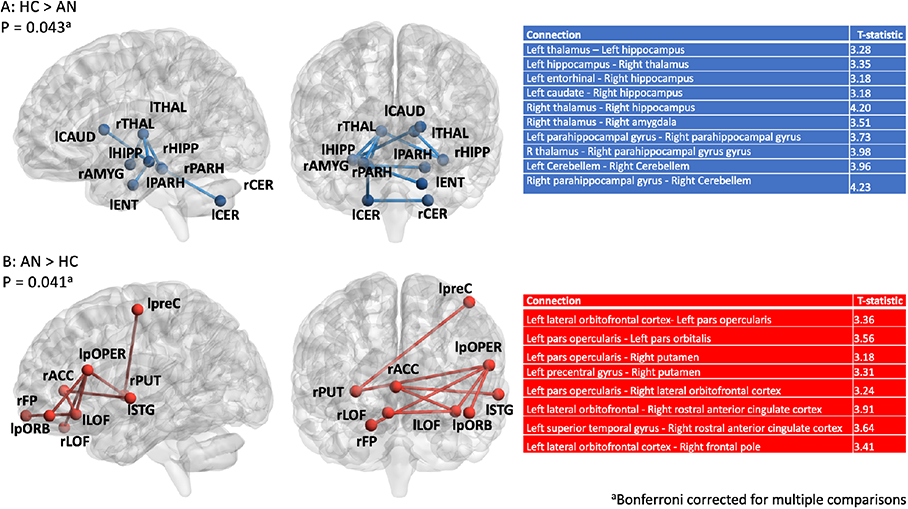
: Weaker subcortical connectivity and stronger cortical connectivity among individuals with AN Panel A) Component comprising tracts with higher streamline counts in HC compared to patients with AN. Panel B) Component comprising tracts with higher streamline counts in patients with AN compared to HC. AN: anorexia nervosa, HC: healthy controls, lLOF: left lateral orbitofrontal gyrus, lpOPER: left pars opercularis, lpORB: left pars orbitalis, lpreC: left precentral gyrus, lSTG: left superior temporal gyrus’ rFP: right frontal pole, rPUT: right putamen, rrACC: right rostral anterior cingulate gyrus, lCer: left Cerebellum, lFP: left frontal pole, lLOF: left lateral orbitofrontal, lPARH: left parahippocampal gyrus, lTHAL: left thalamus, rCer: right Cerebellum, rHIPP: right hippocampus, rPARH: right parahippocampal gyrus, rTHAL: right thalamus. ^a^ P-values are Bonferonni-corrected for multiple comparisons. Node-based analyses

Node-based analyses indicated that individuals with AN, relative to HC, had reduced strength of the left hippocampus (B = -2355.24, 95% CI: -3541.32,-1169.16, p = 0.029). There were no significant differences in local efficiency between groups. Global efficiency did not differ between groups (B = 2.63, 95% CI: [-10.98, 17.31], p = 0.661). Results of all node-based analyses are reported in the Supplement (**Tables S4 and S5**).

### Unharmonized data analyses

Analyses with unharmonized data yielded outcomes that were largely consistent (in direction, size and significance) with outcomes of harmonized data analyses. Full results are reported in the Supplement (**Tables S7-S9**).

### Association between structural connectome metrics and illness severity

Amongst individuals with AN, the standardized BMI variable was positively associated with connectivity within the subcortical component (B = 0.20, 95% CI: [0.04, 0.36], p = 0.016; Figure 3). Illness duration was not significantly associated with any structural connectivity measure.

**Figure 3.**
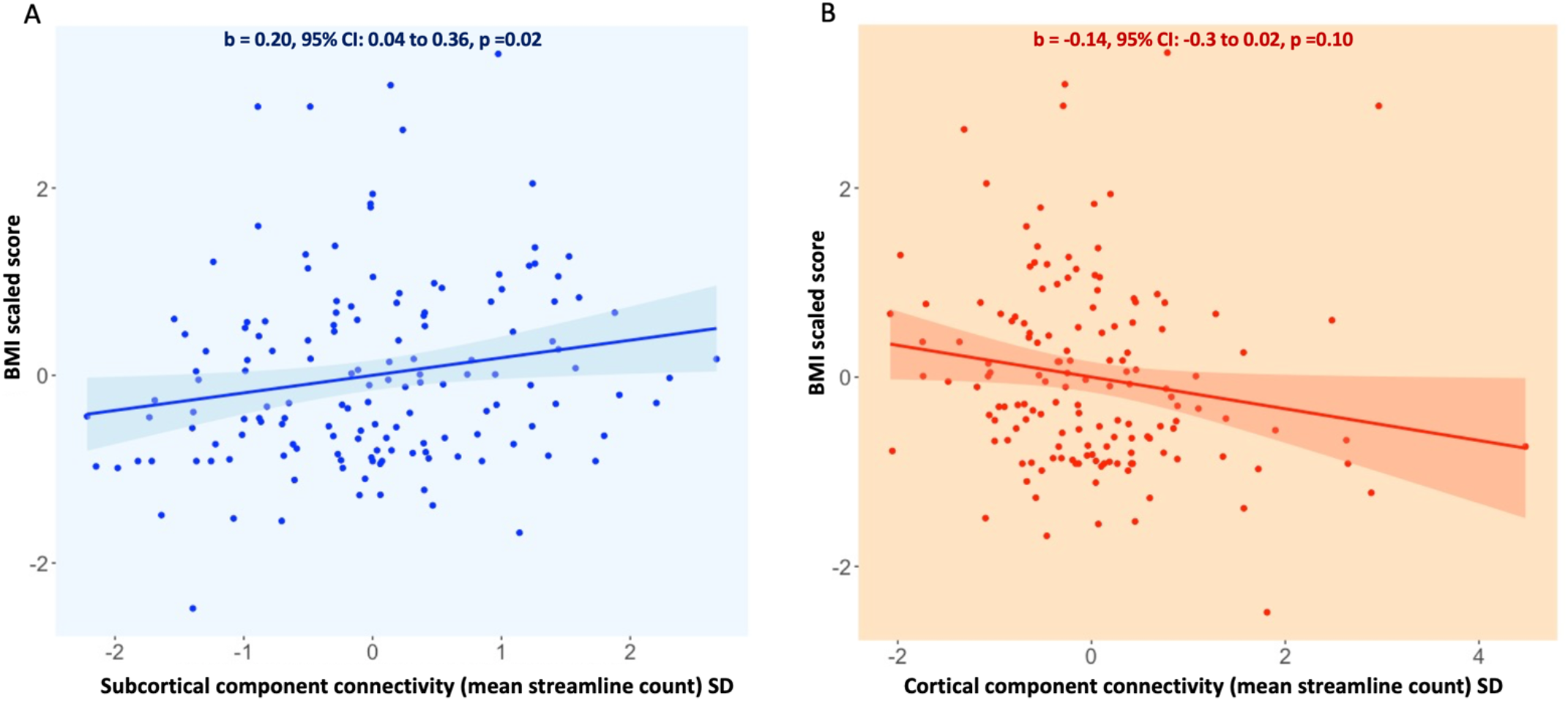
: Associations between subnetwork connectivity and BMI amongst individuals with AN Panel A) Average streamline count in the subcortical network plotted against scaled BMI score in the AN sample. Panel B) Average streamline count in the cortical network plotted against scaled BMI score in the AN sample. The beta value indexes the strength of association between subnetwork connectivity and scaled BMI score, and was calculated by regressing the BMI variable onto the average streamline count within each subnetwork in a model adjusted for age, motion and study. Average streamline counts for each subnetwork were standardized (demeaned and scaled) within the AN group. For the association between subcortical network connectivity and BMI: B = 0.20, 95% CI: 0.04, 0.36, p = 0.02. For the association between cortical network connectivity and BMI: B = 0.14, 95% CI: -0.30, 0.02, p = 0.10.

Full results of these exploratory analyses are presented in **Table 2**.

**Table 2:** Outcomes of linear regression analyses exploring associations of illness severity markers with identified structural connectome differences amongst individuals with AN

| Predictor variable<sup>a</sup> | Association with BMI(kg/m<sup>2</sup>)<sup>bd</sup> |  | Association with Illness Duration<sup>c</sup> |  |
| --- | --- | --- | --- | --- |
|  | B [95% CI] | P-value | B [95% CI] | P-value |
| Structural connectivity metric |  |  |  |  |
| Subcortical component connectivity | 0.2 [0.04,0.36] | 0.02 | -0.02 [-0.14,0.1] | 0.69 |
| Cortical component connectivity | -0.14 [-0.3,0.02] | 0.10 | 0.01 [-0.11,0.13] | 0.86 |
| Strength of left hippocampus | 0.1 [-0.06,0.26] | 0.22 | -0.05 [-0.17,0.07] | 0.40 |
| BMI: body mass index (assessed at the time of scan). |  |  |  |  |
| <sup>a</sup>Predictors and outcome variables were mean centered and scaled. |  |  |  |  |
| <sup>b</sup>Models adjusted for age, motion and study. |  |  |  |  |
| <sup>c</sup>Models adjusted for age, motion, study, and the scaled BMI variable. |  |  |  |  |
| <sup>d</sup>The BMI variable combined scaled BMI scores calculated in adults of the sample (aged over 19 years) with scaled BMI percentile scores calculated in adolescents (aged 19 years and below). |  |  |  |  |

### Subtype differences

AN subtypes did not differ in the connectivity of components or nodes that showed between-group differences in the main analyses (Supplementary material, **Table S10**).

### Subsample analyses

Similar effect estimates for between-group differences in structural connectivity emerged in analyses including only adolescent or only adult participants (Supplementary material, **Table S11**).

## Discussion

The current study used state-of-the-art methods to characterize white matter connectivity of the brain in a large group of individuals with AN, as compared with HC. Anatomical connections of the brain were reconstructed using probabilistic tractography, to recover biologically plausible representations. Structural connectivity data were harmonized using well-validated methods that remove scanner-related variance.

A network-based statistic analysis indicated that, relative to HC, individuals with AN had enhanced connectivity in a cluster of tracts connecting regions of the frontal cortex. In contrast, patients showed reduced connectivity within a network of subcortical brain regions relative to HC. Lower connectivity in this network was associated with lower BMI amongst the AN sample. Node-based analyses identified that the left hippocampus, a region of the subcortical network component that differed between groups, showed weaker connectivity with all regions of the brain amongst individuals with AN relative to HC. These data-driven, whole-brain findings suggest differences in brain organization between individuals with AN and HC that could explain core features of illness. Outcomes of exploratory analyses indicated similar between-group differences in adolescent and adult populations, and no difference in structural connectivity between AN subtypes, suggesting a common set of structural brain alterations across patient subgroups.

The subcortical network that showed alterations in AN overlaps with reward and feeding circuits, and included connections between thalamic, hippocampal, striatal and amygdala regions. Reduced connectivity within this system could impact the desire to eat, as proposed by “bottom-up” models of illness, and may be reflected in the association between subcortical network connectivity and BMI amongst the AN group. Genetic variants favoring low body weight and associated metabolic factors predict increased vulnerability for AN (48); reduced appetitive motivation could mediate genetic influences on body weight and AN development. The amygdala, thalamus, and hippocampus (i.e., regions of the subcortical network that differentiated individuals with AN from HC) are also implicated in anxiety and emotion regulation (49, 50). Abnormalities in connectivity between these regions could contribute to anxiety surrounding eating in AN (51), as well the high levels of general anxiety in AN populations (52).

The heightened frontal connectivity observed amongst individuals with AN is consistent with “top-down” models of illness. These models emphasize greater functioning within frontal circuits (53, 54), which is suggested to underlie the high cognitive control and behavioral rigidity observed in patients (55). The current findings support the possibility that persistent dietary restriction in AN stems from brain network alterations via a combination of reduced appetitive motivation, and inflexible cognition/behavior surrounding eating. This hypothesis can be tested by examining whether the identified network-level structural changes are related to eating behavior.

Several regions of the subcortical network that showed reduced connectivity amongst individuals with AN (i.e., caudate, thalamus, hippocampus) are considered “hubs”, or regions that are typically densely connected. Hubs are particularly well-connected with each other, causing the so-called “rich-club” structure of the brain that ensures a system able to balance segregation and integration for optimal information processing (56). The hippocampus also showed reduced whole-brain connectivity in node-based analyses. The collection of results from this investigation may reflect alterations in global properties of brain organization in AN that hinder adaptive cognition and behavior generally – for example contributing to general psychopathology (e.g., anxiety, depression) or limiting cognitive function (e.g., memory and learning). Two smaller studies have indicated alterations in global brain organization in AN (23, 24), and replicating this work in a larger sample comprises a valuable direction for future research. Disrupted rich club organization has been observed in multiple psychiatric disorders, including major depressive disorder (57), OCD (58) and schizophrenia (59). Similar alterations in AN could explain psychiatric comorbidities of the illness. Notably, node/subnetwork-based analyses of OCD, major depressive disorder, and anxiety disorders have not identified differences between patient and HC populations that match those reported in this study (e.g., (60-62)).

Findings of this study are consistent with prior dMRI studies of white matter microstructure in AN. Several studies have reported that individuals with AN, as compared with HC, have greater integrity within frontal portions of the inferior fronto-occipital fasciculus (63, 64), and reduced integrity of white matter in the fornix (11). The inferior fronto-occipital fasciculus traverses the inferior frontal and medial prefrontal regions in which elevated connectivity was seen amongst individuals with AN in the current study. The fornix comprises the major output tract of the hippocampus, and encapsulates connections of the subcortical component showing reduced connectivity in AN (e.g., tracts between hippocampus, thalamus, parahippocampal cortex and entorhinal cortex). Studies of white matter microstructure do not allow for inferences concerning connectivity between brain regions in AN. However, outcomes from these studies are consistent with greater processing within frontal networks, and reduced processing in subcortical networks, in AN. Although the fornix alterations reported in studies of white matter microstructure are heavily influenced by the ventricular enlargement that occurs with starvation (65), they do not appear to be fully explained by this enlargement (66).

The current findings differ somewhat from prior investigations of region-to-region white matter connectivity in AN. Two prior studies testing group differences in individual tracts reported stronger connectivity in intra-hemispheric frontostriatal tracts amongst individuals with AN, both between the nucleus accumbens and OFC (12, 67), and between the putamen and motor areas (13). These differences were not found in the current study, potentially due to the unit of analysis here being constellations of connected white matter tracts rather than individual tracts. The only other large-scale investigation of the structural connectome in AN probed (as we did) differences in constellations of white matter connections (or subnetworks). This earlier study reported no between-group differences in network connectivity based on streamline counts, but did observe greater integrity of white matter in occipital-parietal regions amongst individuals with AN (14). The tract reconstruction method of the prior study was limited by assuming a single fiber direction per voxel, resulting in a less biologically accurate reconstruction of white matter pathways relative to the multi-fiber probabilistic approach of the current study (15-17). Three smaller studies assessed brain-wide structural connectivity and organization in AN using graph-based approaches (23-25). The only study using probabilistic tractography to recover white matter connections also assessed node-connectivity (23). Inferences from this study (23), including reduced connectivity of frontal brain regions in AN, are largely inconsistent with outcomes of the current investigation. Replication of the tract reconstruction and analytical methods implemented here, ideally within a larger sample, are warranted to confirm the reliability of findings.

This study comprises the largest diffusion tractography investigation of AN to date, and used current best practices for dMRI data processing and data analysis. Nonetheless, there are several limitations that should be considered when interpreting results. First, data are cross-sectional, meaning causes and consequences of starvation cannot be parsed. Longitudinal studies may provide further insight into whether the structural connectivity abnormalities associated with AN in this study comprise mechanisms of illness. It is possible white matter connectivity changes result from the grey matter loss that occurs with low-weight in AN (23, 68, 69). This would not undermine the current findings, but it does further encourage determining whether structural connectivity alterations influence illness onset or maintenance as opposed to constituting illness correlates. Second, ideally dMRI scans would have been acquired with higher b-values (e.g., 3000), for more precise estimation of fiber orientation distributions within a given voxel. However even at low b-values, constrained spherical deconvolution-based tractography outperforms methods assuming a single-fiber orientation within each voxel, in terms of producing an anatomically accurate reconstruction of white matter pathways (70). Third, data harmonization does not completely remove batch effects, and collecting all dMRI data in one study would have been ideal. Though analyses were adjusted for effects of age, it is possible findings are affected by participant groups varying in pubertal stage. Furthermore, statistical analyses could not be adjusted for the effects of handedness as this information was not available for all participants, and whether the lateralization of structural connectivity differs between patients and HC was not explored. Diagnostic assessment and fMRI scanning did not occur on the same day, which introduces a small change in BMI (see Table 1), and some patients initiated treatment (and weight gain) during this time. The time difference was not included as a covariate in analyses as it was small and few patients were affected. It is unlikely this impacted the main conclusions, yet may be considered as a limitation. Finally, the study included females only. AN is more common in females compared to males (71), however whether the neural underpinnings of illness differ by biological sex requires clarification.

In summary, this large-scale investigation of whole-brain structural connectivity in patients with AN implemented advances in diffusion tractography and graph theory methods to yield information about the neural architecture underlying AN. Individuals with AN had reduced connectivity of and between subcortical regions, and elevated connectivity between frontal regions. The identified connectivity profile hints at weaker processing within networks supporting feeding behavior and emotion regulation, and greater processing within networks supporting executive function. This combination may drive restrictive eating behavior, opening new avenues for hypothesis-driven longitudinal research in AN.

## Disclosures

Dr. Steinglass reports receiving royalties from UpTo Date. Dr. Posner has received support from Takeda (formerly Shire), Aevi Genomics, and Innovation Sciences. Dr Lloyd, Dr Foerde, Ms Aw, Mr Semanek, and Dr Muratore have no conflicts of interest to report.

The article has been posted on the bioRxiv preprint server.

