## Supplementary material for "Large-scale exploration of whole-brain structural connectivity in anorexia nervosa: alterations in the connectivity of frontal and subcortical networks"

### Supplementary Materials

#### MRI Scan Parameters

Table S1: MRI scan parameters across the five studies

| Study | Scanner model | T1 structural MRI scan parameters | Diffusion-weighted MRI scan parameters |
|---|---|---|---|
| Study 1 | 1.5 T Philips Intera MRI system with 8-channel head coil | Sequence: T1weighted 3D spoiled gradient recall (SPGR)<br>TR: 2.5 ms<br>TE: 3.7 ms<br>flip angle: 30°<br>FOV: 256 mm<br>128 slices<br>voxel size: 1 × 1 × 1 mm | Diffusion series: two images with minimal diffusion weighting, 16 images with diffusion weighting along non-collinear directions (b-value=800 sm−2)<br>Runs: 2<br>TR: 10,58.6 ms<br>TE:7.0 ms<br>70 slices<br>voxel size: 2 × 2 × 2 mm |
| Study 2 | 3.0T Phillips MRI system with 8-channel head coil. | Sequence: T1weighted Magnetization Prepared Rapid Acquisition Gradient Echo (MP-RAGE)<br>TR: 6.5 ms<br>TE: 3.0 ms<br>flip angle: 8°<br>FOV: 256 mm<br>165 slices<br>voxel size: 1 × 1 × 1 mm | Diffusion series: one image with minimal diffusion weighting, 56 images with diffusion weighting along non-collinear directions (b-value=800 $sm^{-2}$)<br>Runs: 1<br>TR: 775.5 ms<br>TE: 6.8 ms<br>75 slices<br>voxel size: 2 × 2 × 2 mm |
| Study 3 | GE Signa 3T MRI system with 32-channel head coil | Sequence: T1weighted 3D (SPGR) – BRAVO sequence<br>flip angle: 11°<br>FOV: 230 mm<br>voxel size: 0.9 × 0.9 × 0.9 | Diffusion series: three images with minimal diffusion weighting, 25 images with diffusion weighting along non-collinear directions (b-value=1000 sm−2)<br>Runs: 1<br>TR: 850 ms<br>TE: 8.23 ms<br>75 slices<br>voxel size: 0.94 × 0.94 × 2.5 mm |
| Study 4 | GE Signa 3T MRI system with 32-channel head coil | Sequence: T1weighted 3D (SPGR) – BRAVO sequence<br>flip angle: 12°<br>FOV: 256 mm<br>voxel size: 1mm × 1mm × 1mm | Diffusion series: two images with minimal diffusion weighting, 56 images with diffusion weighting along non-collinear directions (b-value=800 $sm^{-2}$)<br>TR: 900 ms<br>TE: minimum<br>75 slices<br>voxel size: 1.5 x 1.5 x 2.0 mm |
| Study 5 scanner 1 | GE Signa 3T MRI system with 32-channel head coil | Sequence: T1weighted 3D (SPGR) – BRAVO sequence<br>flip angle: 12°<br>FOV: 256 mm<br>voxel size: 1mm × 1mm × 1mm | Diffusion series: two images with minimal diffusion weighting, 56 images with diffusion weighting along non-collinear directions (b-value=800 sm−2)<br>TR: 900 ms<br>TE: minimum<br>75 slices<br>Runs: 1<br>voxel size: 1.5 x 1.5 x 2.0 mm |
| Study 5 scanner 2 | GE Signa 3T MRI system with 48-channel head coil | Sequence: T1weighted 3D (SPGR) – BRAVO sequence<br>flip angle: 12°<br>FOV: 256 mm<br>voxel size: 1mm × 1mm × 1mm | Diffusion series: two images with minimal diffusion weighting, 56 images with diffusion weighting along non-collinear directions (b-value=800 $sm^{-2}$)<br>Runs: 1<br>TR: 900 ms<br>TE: minimum<br>75 slices<br>voxel size: 1.5 x 1.5 x 2.0 mm |

**Participant characteristics by study and diagnosis**

Table S2: Participant characteristics by study and diagnosis

| | Study 1 | | Study 2 | | Study 3 | | Study 4 | | Study 5 | |
|---|---|---|---|---|---|---|---|---|---|---|
| | HC (N=20) | AN (N=23) | HC (N=25) | AN (N=26) | HC (N=13) | AN (N=16) | HC (N=35) | AN (N=30) | HC (N=26) | AN (N=52) |
| | Mean ± SD (range) | Mean ± SD (range) | Mean ± SD (range) | Mean ± SD (range) | Mean ± SD (range) | Mean ± SD (range) | Mean ± SD (range) | Mean ± SD (range) | Mean ± SD (range) | Mean ± SD (range) |
| Age (years) | 20.7 ± 2.98 (16, 25) | 19.26 ± 2.49 (16, 25) | 19.36 ± 3.01 (15, 24) | 19.35 ± 3.49 (14, 26) | 22.08 ± 2.47 (18,2 7) | 26.88 ± 6.93 (16, 39) | 25.86 ± 5.35 (19, 39) | 26.93 ± 6.07 (17, 40) | 15.88 ± 1.58 (12, 18) | 15.38 ± 1.62 (12, 18) |
| BMI ($kg/m^2$)[a] | 21.47 ± 1.67 (19.9, 24.3) (N=13) | 17.36 ± 0.92 (15.65, 18.24) (N=8) | 21.24 ± 1.61 (18.9, 23.4) (N=9) | 15.46 ± 2.25 (12.51, 18.12) (N=13) | 21.51 ± 2.33 (19.1, 24.9) (N=11) | 15.32 ± 1.66 (11.72, 17.74) (N=14) | 21 ± 1.60 (18.7, 25.0) (N=32) | 16.00 ± 1.65 (13.52, 18.73) (N=25) | NA (N=0) | NA (N=0) |
| BMI percentile (%)[b] | 35.26 ± 16.7 (19.4, 70.1) (N=7) | 3.92 ± 7.42 (0, 28.6) (N=15) | 50.72 ± 19.23 (21.1, 86.2) (N=16) | 7.64± 6.77 (0, 21.2) (N=13) | 17.75 ± 16.48 (6.1, 29.4) (N=2) | 7.65 ± 3.32 (5.3, 10.0) (N=2) | 22.13 ± 5.45 (16.3, 27.4) (N=3) | 7.62 ± 7.11 (0.7, 17.8) (N=5) | 54.54 ± 24.67 (13.3, 87.4) (N=26) | 9.56 ± 10.46 (0, 39.6) (N=52) |
| EDEQ Global score | 0.09 ± 0.14 (0, 0.45) | 3.99 ± 1.36 (1.46, 5.65) | 0.19 ± 0.2 (0, 0.75) | 4.14 ± 1.41 (0.30, 5.6) | 0.09 ± 0.09 (0, 0.25) | 4.43 ± 1.14 (1.38, 5.65) | 0.35 ± 0.48 (0, 0.2.09) | 4.27 ± 1.5 (0.03, 5.80) | 0.42 ± 0.49 (0, 0.2.17) | 3.39 ± 1.68 (0, 6) |
| Illness Duration (months) | NA ± NA | 52.48 ± 37.72 (3, 120) | NA ± NA | 38.19 ± 28.42 (8, 120) | NA ± NA | 144.5 ± 82.34 (12, 278) | NA ± NA | 118.53 ± 79.39 (12, 324) | NA ± NA | 8.51 ± 11.8 (0, 59) |
| Mean framewise displacement across DWI series | 0.2 ± 0.1 | 0.26 ± 0.16 | 0.11 ± 0.13 | 0.05 ± 0.07 | 0.32 ± 0.04 | 0.33 ± 0.03 | 0.12 ± 0.05 | 0.11 ± 0.04 | 0.16 ± 0.07 | 0.2 ± 0.17 |

[a] BMI information presented for participants over 19
[b] BMI percentile information presented for participants 19 and under
BMI: Body mass index; EDEQ: Eating disorder examination questionnaire

#### Methodological details

*Probabilistic tractography*

Response functions of white matter, grey matter and CSF were derived with the dhollander algorithm (1), using the dMRI signal and the five-tissue type image generated from the structural MRI scan. Next, the response functions of white matter fibres were estimated in the voxels with the 300 greatest fractional anisotropy values (indicative of more directed diffusion). Using the dMRI signal and the calculated response functions, the underlying distribution of fibers in each voxel was recovered using multi-shell multi-tissue constrained spherical deconvolution (2), with default maximum spherical harmonic degree parameters. This approach allows dMRI signal resulting from white matter and CSF components to be distinguished, preventing CSF contaminating estimates of white matter fiber orientation distributions. White matter fiber orientation distributions were normalized, to remove residual inhomogeneity in intensities. Seed points for streamlines were determined dynamically. Seeding continued until 20,000,000 streamlines between 10 and 250 mm were generated. Streamline step size was 0.625 mm; the maximum angle between successive steps was 22.5 degrees.

The iFOD2 (3) tracking algorithm determines the probability of candidate streamline paths by sampling fiber orientation distributions along the entire path. More probable, and thus reconstructed, streamlines are generally those where fiber orientation distribution amplitudes along the path are large. The tractography was anatomically constrained, with the five-tissue type image used as a biological prior to guide streamline termination and acceptance (4). Streamlines were re-tracked if a poor termination was encountered, and cropped at the grey matter white matter interface.

*Network-based statistic*

To complete the network-based statistic analysis, first tracts differing in streamline count between groups at a t-score threshold of greater than 3.1 (corresponding to a p-value of 0.001) are identified. Next, components, or sets of connected tracts meeting the t-score criteria, are identified. The null distribution of the maximum component size is empirically estimated using a permutation approach. P-values for the hypothesis that observed components (i.e., those identified in the unpermuted data) are larger than expected should there be no between-group difference in subnetwork connectivity can then be derived. 10,000 permutations were specified, and identified components were significant at a family-wise error corrected rate of $p < 0.05$.

*Node-based analyses*

Strength is calculated as the total number of connections (streamlines) of each node. To calculate efficiency, the inverse of the shortest path length between all nodes directly connected to the node in question was determined and averaged. Path length was defined as the inverse of streamline count. The contribution of a given path to efficiency calculations was proportionate to the streamline count between the path nodes and the node for which efficiency was calculated (i.e., the average was weighted).

#### Details of component connectivity by study and diagnosis

Table S3: Connectivity in subcortical and cortical components by study and diagnosis

| | Study 1 | | Study 2 | | Study 3 | | Study 4 | | Study 5 | | Full sample | |
|---|---|---|---|---|---|---|---|---|---|---|---|---|
| | HC (N=20) | AN (N=23) | HC (N=25) | AN (N=26) | HC (N=13) | AN (N=16) | HC (N=35) | AN (N=30) | HC (N=26) | AN (N=52) | HC (N=119) | AN (N=147) |
| | Mean (SD) | Mean (SD) | Mean (SD) | Mean (SD) | Mean (SD) | Mean (SD) | Mean (SD) | Mean (SD) | Mean (SD) | Mean (SD) | Mean (SD) | Mean (SD) |
| Subcortical component connectivity | 6416.44 (1044.80) | 5926.73 (779.30) | 6490.86 (933.36) | 5937.17 (751.08) | 6775.51 (1710.19) | 5481.15 (677.18) | 6387.86 (832.13) | 5637.40 (804.14) | 6536.10 (793.97) | 5741.32 (832.15) | 6849.04 (998.46) | 5755.44 (791.98) |
| Cortical component connectivity | 131.27 (33.42) | 202.50 (70.40) | 155.28 (66.00) | 186.54 (49.70) | 145.03 (72.41) | 209.88 (67.63) | 164.51 (62.49) | 188.10 (71.11) | 133.12 (45.27) | 182.76 (64.36) | 148.00 (57.86) | 190.56 (64.66) |

Connectivity defined as average streamline count in component connections

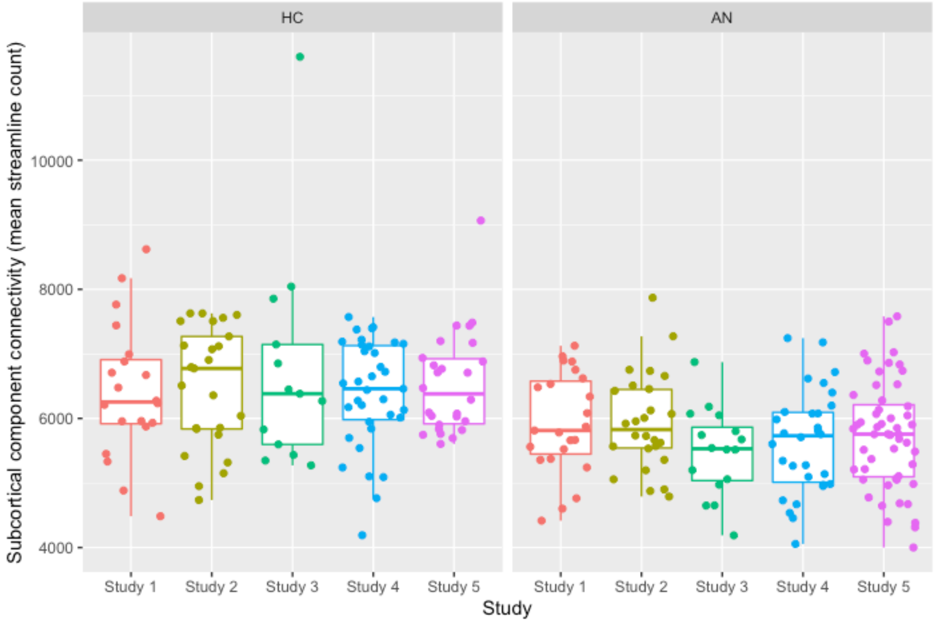

Figure S1: Distribution of subcortical connectivity values across studies and diagnostic groups

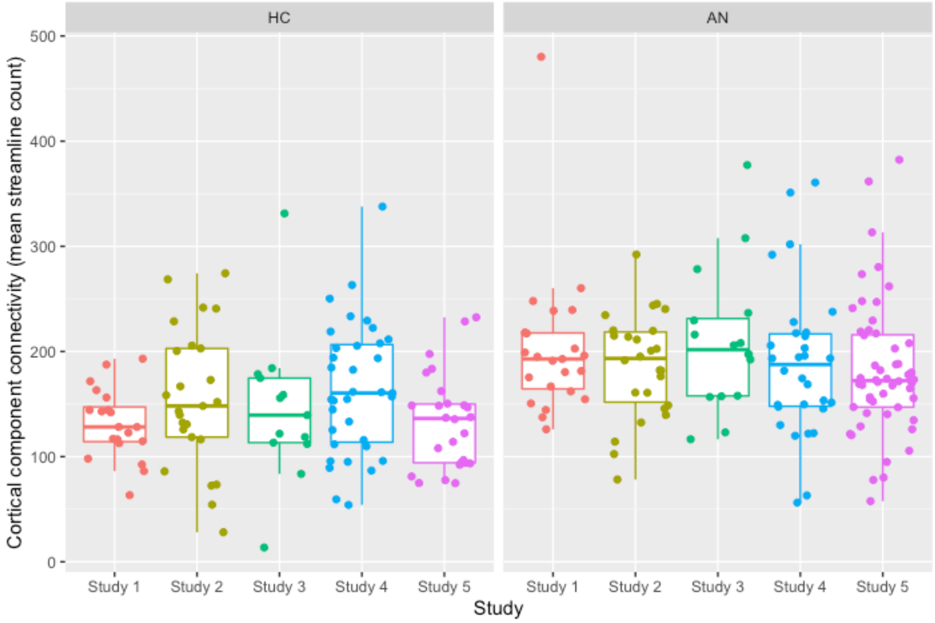

Figure S2: Distribution of cortical connectivity values across studies and diagnostic groups

**Full results of node analyses**

Table S4: Outcomes of between-group analyses of node strength

| | Strength | | | |
|---|---|---|---|---|
| Node | HC | AN | Effect of Diagnosis | |
| | Mean (SD) | Mean (SD) | B [95% CI] | P value |
| L nucleus accumbens | 18389.17 (4198.67) | 17974.38 (4067.03) | -311.28 [-802.51,179.94] | 1.00 |
| L amygdala | 27749.78 (5904.56) | 26374.12 (4220.97) | -701.5 [-1323.83,-79.18] | 0.94 |
| L bank of the superior temporal sulcu s | 27906.4 (5144.26) | 28991.09 (5486.11) | 319.03 [-498.78,1136.84] | 1.00 |
| L caudal anterior cingulate | 99364.08 (15726.24) | 97915.09 (13662.45) | 553.4 [-106.99,1213.79] | 1.00 |
| L caudate | 50326.11 (10737.9) | 45392.74 (9210.79) | -839.54 [-2637.6,958.52] | 1.00 |
| L Cerebellum | 64414.45 (12615.79) | 65527.85 (9412.9) | -2395.72 [-3623.23,-1168.22] | 0.06 |
| L caudal middle frontal gyrus | 42232.52 (8374.17) | 43018.3 (7785.65) | 361.43 [-959.77,1682.64] | 1.00 |
| L cuneus | 21899.06 (5752.99) | 20386.96 (5530.44) | 373.33 [-624.59,1371.25] | 1.00 |
| L entorhinal | 7774.54 (2825.44) | 8305.6 (3155.26) | -692.35 [-1382.38,-2.31] | 0.99 |
| L frontal pole | 63654.04 (9810.32) | 65686.99 (9113.06) | 328.44 [-28.31,685.19] | 1.00 |
| L fusiform | 80666.09 (10483.03) | 75969.65 (8917.33) | 1039.19 [-127.7,2206.07] | 1.00 |
| L hippocampus | 39019.78 (4848.8) | 39767.21 (5136.66) | -2355.24 [-3541.32,-1169.16] | **0.03** |
| L isthmus cingulate cortex | 102919.25 (13855.57) | 103540.69 (13199.9) | 399.36 [-221.75,1020.48] | 1.00 |
| L insula | 85450.19 (12367.1) | 87898.53 (11749.97) | 205 [-1458.06,1868.07] | 1.00 |
| L inferior parietal lobule | 65169.68 (13043.82) | 68417.04 (11012.67) | 1232.85 [-253.6,2719.31] | 1.00 |
| L inferior temporal gyrus | 54732.06 (9355.99) | 54006.02 (8059.07) | 1594.46 [130.92,3058] | 0.98 |
| L lingual | 65186.75 (10166.63) | 65671.89 (9204.88) | -310.67 [-1374.4,753.05] | 1.00 |
| L lateral orbitofrontal | 81573.36 (14175.5) | 85720.1 (12204.93) | 412.34 [-742.74,1567.41] | 1.00 |
| L lateral occipital gyrus | 55233.55 (11103.85) | 53678.09 (10448.94) | 2084.49 [465.38,3703.59] | 0.64 |
| L medial orbitofrontal | 64818.41 (11594.07) | 66195.92 (10276.11) | -586.75 [-1858.92,685.41] | 1.00 |
| L middle temporal gyrus | 53561.93 (9282.26) | 52203.42 (9358.82) | 781.76 [-550.9,2114.42] | 1.00 |
| L pallidum | 45527.75 (7555.4) | 45552.25 (6237.98) | -637.38 [-1781.45,506.69] | 1.00 |
| L paracentral | 34559.35 (6502.3) | 32038.22 (5001.63) | 14.79 [-834.75,864.33] | 1.00 |
| L parahippocampal | 38140.35 (6238.79) | 39152.37 (5686.65) | -1254.28 [-1962.52,-546.04] | 0.16 |
| L posterior cingulate cortex | 78972.58 (10704.07) | 80334.9 (9018.85) | 527.69 [-207.9,1263.28] | 1.00 |
| L precuneus | 57726.65 (12722.2) | 57813.28 (12166.32) | 687.75 [-522.22,1897.71] | 1.00 |
| L pericalcarine | 44179.22 (8890.28) | 46366.89 (7572.4) | 57.89 [-1480.94,1596.72] | 1.00 |
| L pars opercularis | 21814.5 (4641.77) | 23041.06 (4122.26) | 1053.46 [40.35,2066.56] | 0.99 |
| L pars orbitalis | 79455.85 (13276.01) | 81279.89 (9722.93) | 667.73 [139.83,1195.63] | 0.79 |
| L postcentral | 120601.68 (17788.59) | 123136.37 (10851.74) | 897.87 [-508.07,2303.8] | 1.00 |
| L precentral | 37786.95 (6877.09) | 38650.06 (6283.18) | 1278.34 [-484.97,3041.66] | 1.00 |
| L pars triangularis | 131869.5 (15898.46) | 134289.9 (17003.15) | 504.31 [-296.21,1304.83] | 1.00 |
| L putamen | 33277.51 (5613.13) | 32354.78 (6076.02) | 1160.98 [-886,3207.97] | 1.00 |
| L rostral anterior cingulate cortex | 108012.94 (20363.75) | 109460.38 (16610.92) | -390.91 [-1113.38,331.55] | 1.00 |
| L rostral middle frontal gyrus | 148423.79 (22415.09) | 151976.54 (12962.65) | 520.86 [-1693.49,2735.2] | 1.00 |
| L superior frontal gyrus | 74717.74 (12189.97) | 75008.9 (9269.08) | 1871.64 [-309.67,4052.94] | 1.00 |
| L supramarginal gyrus | 115783.09 (16917.19) | 120692.21 (14209.67) | 219.27 [-1085.22,1523.76] | 1.00 |

| | | | | |
|---|---|---|---|---|
| L superior parietal lobule | 65110.31 (9887.31) | 66699.57 (7663.55) | 2419.54 [502.02,4337.07] | 0.75 |
| L superior temporal gyrus | 121004.22 (12808.14) | 118412.9 (12493.77) | 873.03 [-181.7,1927.75] | 1.00 |
| L thalamus | 13019.88 (3838.87) | 13361.03 (3825.2) | -1262.04 [-2824.18,300.1] | 1.00 |
| L temporal pole | 15261.47 (2640.69) | 15371.1 (2645.78) | 224.18 [-242.7,691.07] | 1.00 |
| L transverse temporal | 20868.93 (5265.57) | 20675.17 (4720.58) | 62.04 [-264.97,389.05] | 1.00 |
| R nucleus accumbens | 31892.98 (6450.73) | 31072.06 (5572.3) | -171.09 [-768.28,426.09] | 1.00 |
| R amygdala | 32237.61 (7672.41) | 32444.17 (7190.9) | -415.16 [-1151.63,321.31] | 1.00 |
| R bank of the superior temporal sulc us | 30873.63 (6367.49) | 31918.59 (5336.83) | 8.13 [-906.17,922.43] | 1.00 |
| R caudal anterior cingulate | 106170.67 (17107.12) | 102932.17 (13602.37) | 519.6 [-200.64,1239.84] | 1.00 |
| R caudate | 49935.46 (11374.47) | 45116.28 (9144.79) | -1699.2 [-3567.05,168.65] | 1.00 |
| R Cerebellum | 63324 (11228.14) | 64026.49 (9839.43) | -2346.37 [-3607.42,-1085.32] | 0.08 |
| R caudal middle frontal gyrus | 45187.61 (7423.23) | 45031.97 (6544.54) | 211.1 [-1080.71,1502.9] | 1.00 |
| R cuneus | 22180.32 (7083.62) | 20729.37 (5786.77) | -84.96 [-946.47,776.55] | 1.00 |
| R entorhinal | 9829.98 (3582.88) | 10536.21 (3641.89) | -691.35 [-1482.87,100.16] | 1.00 |
| R frontal pole | 65106.13 (10536.38) | 67015.19 (9005.85) | 416.62 [-17.87,851.11] | 1.00 |
| R fusiform | 87179.09 (12306.75) | 82671.57 (11718.9) | 806.36 [-390.03,2002.75] | 1.00 |
| R hippocampus | 35932.18 (5375.78) | 36768.52 (4484.05) | -2305.62 [-3775.03,-836.21] | 0.25 |
| R isthmus cingulate cortex | 106980.78 (18994.13) | 105796.18 (17065.22) | 445.55 [-160.34,1051.44] | 1.00 |
| R insula | 93015.81 (11744.73) | 94052.43 (10003.75) | -691.48 [-2901.4,1518.45] | 1.00 |
| R inferior parietal lobule | 65640.18 (12695.82) | 66573.13 (9903.84) | 417.29 [-912,1746.58] | 1.00 |
| R inferior temporal gyrus | 59333.64 (9108.95) | 58330.36 (7203.8) | 481.94 [-898.41,1862.3] | 1.00 |
| R lingual | 71098.94 (11251.34) | 71108.06 (11217.79) | -470.73 [-1470.24,528.77] | 1.00 |
| R lateral orbitofrontal | 84288.7 (10590.29) | 88984.61 (10395.61) | -3.27 [-1393.76,1387.21] | 1.00 |
| R lateral occipital gyrus | 48282.44 (8692.18) | 48062.14 (8813.03) | 2221.27 [932.73,3509.82] | 0.13 |
| R medial orbitofrontal | 69895.41 (12283.73) | 70355.41 (10301.23) | -36.37 [-1094.57,1021.83] | 1.00 |
| R middle temporal gyrus | 54167.02 (10715.81) | 54762.44 (9892.68) | 358.48 [-1007.64,1724.6] | 1.00 |
| R pallidum | 50028.63 (7548.67) | 49469.08 (5741.22) | 323.55 [-934.7,1581.79] | 1.00 |
| R paracentral | 35641.03 (6563.57) | 33121.44 (5695.5) | -292.12 [-1111.14,526.9] | 1.00 |
| R parahippocampal | 38764.94 (6305.75) | 39503.5 (5790) | -1244.24 [-1995.59,-492.88] | 0.14 |
| R posterior cingulate cortex | 81153.17 (9862.27) | 82135.35 (8854.76) | 399.16 [-345.15,1143.48] | 1.00 |
| R precuneus | 61830.88 (10683.31) | 61520.67 (11118.76) | 320.28 [-819.58,1460.14] | 1.00 |
| R pericalcarine | 42922.5 (7286.31) | 43774.25 (7472.63) | -227.16 [-1579.33,1125.02] | 1.00 |
| R pars opercularis | 25948.65 (5260.2) | 26237.13 (4826.96) | 293.41 [-607.08,1193.91] | 1.00 |
| R pars orbitalis | 80831.79 (12135.72) | 81440.85 (9402.37) | 165.26 [-454.64,785.15] | 1.00 |
| R postcentral | 121139.3 (15534.72) | 122700.89 (10633.16) | 193.31 [-1118.84,1505.46] | 1.00 |
| R precentral | 40403.43 (6248.98) | 40997.79 (7602.87) | 644.69 [-966.07,2255.45] | 1.00 |
| R pars triangularis | 135916.46 (17490.07) | 140070.36 (19319.68) | 316.19 [-552.1,1184.49] | 1.00 |
| R putamen | 27236.12 (5049.53) | 27443.49 (4655.53) | 1835.75 [-431.77,4103.28] | 1.00 |
| R rostral anterior cingulate cortex | 113566.14 (20442.61) | 113761.1 (17211.38) | 131.27 [-468.1,730.65] | 1.00 |
| R rostral middle frontal gyrus | 144200.13 (19088.73) | 147624.88 (13601.67) | -109.14 [-2397.05,2178.78] | 1.00 |
| R superior frontal gyrus | 71723.12 (10958.64) | 72446.73 (9737.84) | 1716.21 [-289.11,3721.53] | 1.00 |
| R supramarginal gyrus | 113413.8 (15354.8) | 118268.35 (13268.01) | 331.24 [-931.3,1593.79] | 1.00 |
| R superior parietal lobule | 60595.97 (8707.41) | 61248.33 (9761.85) | 2233.3 [475.8,3990.81] | 0.72 |
| R superior temporal gyrus | 113115.18 (12392.05) | 108095.55 (10519.94) | 429.43 [-695.35,1554.2] | 1.00 |

| | | | | |
|---|---|---|---|---|
| R thalamus | 14454.45 (5107.53) | 14485.12 (3962.85) | -2416.64 [-3817.04,-1016.23] | 0.09 |
| R temporal pole | 13632.88 (2488.46) | 13309.64 (2202.04) | 47.6 [-506.26,601.46] | 1.00 |
| R transverse temporal | 18389.17 (4198.67) | 17974.38 (4067.03) | -133.81 [-416.27,148.65] | 1.00 |
| R: right, L:left. All models adjusted for age, motion, study. [a] P values permutation-corrected for multiple comparisons. | | | | |

Table S5: Outcomes of between-group analyses of node efficiency

| | Efficiency | | | |
|---|---|---|---|---|
| Node | HC | AN | Effect of Diagnosis | |
| | Mean (SD) | Mean (SD) | B [95% CI] | P value |
| L nucleus accumbens | 108.34 (23.8) | 107.93 (26.45) | -0.74 [-3.81,2.33] | 1.00 |
| L amygdala | 137.77 (24.68) | 133.7 (21.52) | -1.97 [-4.82,0.87] | 1.00 |
| L bank of the superior temporal sulcus | 132.92 (28.63) | 139.28 (27.8) | 3.13 [-0.36,6.62] | 1.00 |
| L caudal anterior cingulate | 542.95 (92.37) | 536.52 (77.5) | 3.71 [-0.27,7.7] | 1.00 |
| L caudate | 185.58 (38.96) | 178.83 (43.05) | -3.75 [-14.18,6.69] | 1.00 |
| L cerebellum | 336.19 (61.18) | 334.19 (44.93) | -2.91 [-7.98,2.16] | 1.00 |
| L caudal middle frontal gyrus | 120.53 (27.55) | 122.65 (27.49) | -0.77 [-7.26,5.72] | 1.00 |
| L cuneus | 68.47 (20.03) | 61.98 (16.93) | 1.06 [-2.35,4.47] | 1.00 |
| L entorhinal | 38.34 (13.44) | 40.74 (15.63) | -3.09 [-5.33,-0.85] | 0.56 |
| L frontal pole | 253.23 (44.84) | 259.69 (45.57) | 1.54 [-0.2,3.28] | 1.00 |
| L fusiform | 378.63 (46.09) | 360.85 (50.6) | 3.31 [-2.29,8.91] | 1.00 |
| L hippocampus | 303.26 (45.99) | 307.04 (46.05) | -8.4 [-14.39,-2.41] | 0.56 |
| L isthmus cingulate cortex | 608 (81.69) | 617.23 (78.25) | 2.15 [-3.53,7.84] | 1.00 |
| L insula | 433.4 (69.49) | 444.78 (59.42) | 4.14 [-5.7,13.97] | 1.00 |
| L inferior parietal lobule | 311.74 (59.87) | 326.72 (51.97) | 6.26 [-1.58,14.1] | 1.00 |
| L inferior temporal gyrus | 188.94 (31.39) | 185.15 (30.89) | 7.58 [0.72,14.44] | 0.96 |
| L lingual | 265.95 (43.05) | 279.31 (38.37) | -1.53 [-5.35,2.28] | 1.00 |
| L lateral orbitofrontal | 312.51 (55.58) | 318.59 (57.49) | 7.6 [2.79,12.42] | 0.23 |
| L lateral occipital gyrus | 291.95 (77.28) | 285 (77.7) | 3.44 [-3.49,10.37] | 1.00 |
| L medial orbitofrontal | 264.73 (48.53) | 272.43 (50.83) | -2.04 [-11.17,7.09] | 1.00 |
| L middle temporal gyrus | 411.84 (69.49) | 409.97 (69.73) | 5.05 [-0.95,11.05] | 1.00 |
| L pallidum | 235.42 (40.27) | 236.15 (32.11) | 0.1 [-8.46,8.65] | 1.00 |
| L paracentral | 146.45 (35.36) | 137.82 (32.21) | 0.72 [-3.68,5.12] | 1.00 |
| L parahippocampal | 276.47 (51.42) | 279.08 (42.94) | -3.95 [-8.1,0.2] | 1.00 |
| L posterior cingulate cortex | 493.3 (71.52) | 494.29 (60.35) | 1.63 [-4.16,7.41] | 1.00 |
| L precuneus | 146.91 (31.55) | 146.88 (34.06) | 1.21 [-6.78,9.2] | 1.00 |
| L pericalcarine | 247.61 (50.83) | 259.63 (38.94) | 0.22 [-3.87,4.3] | 1.00 |
| L pars opercularis | 85.83 (20.97) | 93.82 (18.94) | 6.75 [1.3,12.21] | 0.82 |
| L pars orbitalis | 319.27 (56.3) | 329.63 (43.71) | 4.35 [1.96,6.73] | 0.04 |
| L postcentral | 544.52 (88.12) | 550.62 (60.61) | 5.84 [-0.26,11.94] | 1.00 |
| L precentral | 176.92 (35.63) | 184.24 (28.96) | 4.13 [-4.92,13.18] | 1.00 |
| L pars triangularis | 758.28 (90.37) | 780.56 (89.33) | 4.4 [0.57,8.22] | 0.91 |
| L putamen | 193.09 (45.6) | 188.81 (41.57) | 12.02 [1,23.04] | 0.96 |
| L rostral anterior cingulate cortex | 496.01 (80.16) | 508.25 (62.38) | -1.64 [-6.97,3.69] | 1.00 |
| L rostral middle frontal gyrus | 744.56 (110.72) | 757.29 (79.5) | 6.68 [-1.93,15.29] | 1.00 |
| L superior frontal gyrus | 369.12 (62.46) | 369.43 (52.46) | 7.92 [-3.65,19.49] | 1.00 |
| L supramarginal gyrus | 591.91 (87.75) | 612.48 (71.92) | 0.87 [-6.1,7.84] | 1.00 |
| L superior parietal lobule | 317.55 (50.69) | 324.96 (42.85) | 11.28 [1.59,20.97] | 0.90 |
| L superior temporal gyrus | 904.32 (94.06) | 896.91 (91.78) | 4.39 [-1.23,10.01] | 1.00 |
| L thalamus | 45.66 (13.96) | 46.28 (12.86) | -3.21 [-14.67,8.24] | 1.00 |

| | | | | |
|---|---|---|---|---|
| L temporal pole | 84.49 (17.81) | 84.8 (14.96) | 0.47 [-1.14,2.09] | 1.00 |
| L transverse temporal | 127 (29.79) | 124.47 (34.72) | 0.16 [-1.84,2.17] | 1.00 |
| R nucleus accumbens | 158.63 (26.36) | 153.92 (28.57) | -1.54 [-5.53,2.44] | 1.00 |
| R amygdala | 145.4 (31.33) | 152.3 (31) | -2.29 [-5.69,1.1] | 1.00 |
| R bank of the superior temporal sulcus | 204.01 (37.61) | 211 (33.76) | 3.12 [-0.73,6.98] | 1.00 |
| R caudal anterior cingulate | 592.16 (95.06) | 576.39 (83.85) | 3.76 [-0.63,8.15] | 1.00 |
| R caudate | 181.98 (44.44) | 175 (39.41) | -7.99 [-18.94,2.96] | 1.00 |
| R cerebellum | 329.08 (51.75) | 331.07 (45.95) | -3.05 [-8.18,2.08] | 1.00 |
| R caudal middle frontal gyrus | 125.22 (23.34) | 123.98 (23.35) | 0.91 [-5.09,6.91] | 1.00 |
| R cuneus | 64.31 (21.71) | 60.74 (19.53) | -0.49 [-3.37,2.38] | 1.00 |
| R entorhinal | 43.85 (14.49) | 47.83 (17.66) | -1.57 [-4.06,0.92] | 1.00 |
| R frontal pole | 238.31 (41.92) | 244.59 (43.45) | 2.39 [0.45,4.33] | 0.81 |
| R fusiform | 437.99 (54.61) | 422.71 (58.46) | 2.47 [-2.81,7.75] | 1.00 |
| R hippocampus | 266.9 (47.25) | 272.49 (39.67) | -7.32 [-14.31,-0.33] | 0.99 |
| R isthmus cingulate cortex | 633.25 (102.03) | 616.3 (90.71) | 3.25 [-2.05,8.56] | 1.00 |
| R insula | 485.51 (66.35) | 492.77 (64.71) | -8.5 [-20.27,3.27] | 1.00 |
| R inferior parietal lobule | 290.53 (43.91) | 293.98 (47.75) | 3.91 [-4.13,11.95] | 1.00 |
| R inferior temporal gyrus | 203.41 (33.69) | 197.77 (32.39) | 2.2 [-3.46,7.86] | 1.00 |
| R lingual | 310.03 (66.11) | 325.07 (65.58) | -2.61 [-6.68,1.46] | 1.00 |
| R lateral orbitofrontal | 314.99 (49.67) | 326.18 (50.65) | 7.75 [-0.39,15.9] | 1.00 |
| R lateral occipital gyrus | 220.09 (63.02) | 219.91 (55.8) | 5.87 [-0.27,12.02] | 1.00 |
| R medial orbitofrontal | 285.7 (48.4) | 291.69 (51.92) | 0.2 [-7.03,7.43] | 1.00 |
| R middle temporal gyrus | 408.27 (70.74) | 415.49 (65.06) | 4.44 [-1.53,10.41] | 1.00 |
| R pallidum | 265.02 (37.64) | 264.2 (37.23) | 4.26 [-4.05,12.57] | 1.00 |
| R paracentral | 159.26 (37.48) | 147.25 (35.91) | 0.07 [-4.53,4.67] | 1.00 |
| R parahippocampal | 278.15 (49.32) | 282.27 (42.84) | -5.69 [-10.19,-1.19] | 0.78 |
| R posterior cingulate cortex | 491.76 (61.46) | 492.25 (50.21) | 2.32 [-3.32,7.96] | 1.00 |
| R precuneus | 147.23 (26.73) | 146.21 (29.01) | 0.44 [-6.39,7.27] | 1.00 |
| R pericalcarine | 229.53 (41.75) | 234.66 (37.31) | -0.49 [-3.93,2.96] | 1.00 |
| R pars opercularis | 99.61 (22.91) | 104.68 (24.95) | 2.58 [-2.29,7.45] | 1.00 |
| R pars orbitalis | 328.2 (47) | 325.74 (44.65) | 2.96 [0.04,5.88] | 0.99 |
| R postcentral | 546.07 (70.51) | 547.96 (59.81) | -0.81 [-6.43,4.81] | 1.00 |
| R precentral | 189.55 (36.31) | 195.84 (36.66) | 1.6 [-6.36,9.57] | 1.00 |
| R pars triangularis | 780.61 (97.23) | 807.56 (97.44) | 3.72 [-0.73,8.17] | 1.00 |
| R putamen | 147.41 (23.61) | 149.63 (24.63) | 13.5 [1.43,25.57] | 0.96 |
| R rostral anterior cingulate cortex | 519.56 (76.61) | 529.03 (70.03) | 1.33 [-1.64,4.29] | 1.00 |
| R rostral middle frontal gyrus | 732.98 (98.9) | 747.13 (80.65) | 5.32 [-3.62,14.25] | 1.00 |
| R superior frontal gyrus | 357.59 (53.96) | 354.35 (54.5) | 8.25 [-2.7,19.21] | 1.00 |
| R supramarginal gyrus | 593.27 (78.13) | 609 (65.76) | -1.19 [-7.85,5.47] | 1.00 |
| R superior parietal lobule | 301.57 (51.46) | 306.01 (46.36) | 8.38 [-0.32,17.07] | 1.00 |
| R superior temporal gyrus | 867.27 (91) | 840.48 (83.83) | 2.83 [-3.07,8.73] | 1.00 |
| R thalamus | 45.89 (12.88) | 46.42 (12.41) | -12.91 [-23.63,-2.19] | 0.89 |
| R temporal pole | 78.71 (18.14) | 79.71 (16.32) | 0.43 [-1.11,1.98] | 1.00 |
| R transverse temporal | 108.34 (23.8) | 107.93 (26.45) | 0.65 [-1.44,2.74] | 1.00 |
| R: right, L:left. All models adjusted for age, motion, study. [a] P values permutation-corrected for multiple comparisons | | | | |

Table S6: Node metrics by diagnostic group and study

| | Study 1 | | Study 2 | | Study 3 | | Study 4 | | Study 5 | |
|---|---|---|---|---|---|---|---|---|---|---|
| STRENGTH | HC (N=20) | AN (N=23) | HC (N=25) | AN (N=26) | HC (N=13) | AN (N=16) | HC (N=35) | AN (N=30) | HC (N=26) | AN (N=52) |
| | Mean (SD) | Mean (SD) | Mean (SD) | Mean (SD) | Mean (SD) | Mean (SD) | Mean (SD) | Mean (SD) | Mean (SD) | Mean (SD) |
| L nucleus accumbens | 18120.66 (4985.79) | 18241.2 (3427.16) | 18607.14 (3370.19) | 17480.14 (4277.6) | 17138.18 (4739.15) | 17455.58 (4741.11) | 17583.02 (3164.91) | 16668.47 (3766.61) | 20096.82 (4914) | 19016.51 (4035.56) |
| L amygdala | 27110.76 (4043.22) | 27119.45 (4652.87) | 27858.4 (5870.55) | 25733.41 (4092.42) | 29984.98 (5355.73) | 26274.17 (4930.81) | 26450.65 (5607.27) | 26915.77 (4606.87) | 28768.11 (7492.79) | 26083.08 (3676.26) |
| L bank of the superior temporal sulcus | 30531.89 (6242.96) | 31583.26 (7234.34) | 32071.4 (9448.37) | 30439.31 (4786.48) | 29954.7 (7630.89) | 33211.54 (4565.14) | 30198.17 (6353.54) | 32322.76 (5603.19) | 31870.14 (6004.48) | 31687.99 (7091.65) |
| L caudal anterior cingulate | 27306.84 (7413.89) | 29419.15 (4970.6) | 27998.58 (4504.62) | 28805.8 (4879.98) | 27422.43 (3252.56) | 30349.28 (5328.27) | 27731.77 (5215.98) | 29618.5 (6005.16) | 28756.04 (4512.54) | 28114.54 (5768.88) |
| L caudate | 98916.97 (13947.24) | 100760.67 (12562.6) | 98986.55 (14751.6) | 99580.39 (18610.71) | 94802.77 (20383.3) | 100392.73 (11271.09) | 96425.4 (13562.3) | 95962.12 (11834.76) | 106307.57 (16960.1) | 96188.18 (12931.44) |
| L Cerebellum | 49203.12 (10453.79) | 45719.77 (9340.36) | 50812.53 (11418.39) | 47762.12 (9278.44) | 51523.54 (15751.79) | 44918.23 (8315.8) | 51152.35 (9735.77) | 43835.94 (8210.36) | 49011.29 (9124.67) | 45107.56 (9993.86) |
| L caudal middle frontal gyrus | 61435.17 (15865.95) | 68056.38 (4557.02) | 67179.27 (14257.95) | 61041.31 (8863.67) | 63481.96 (19018.48) | 68597.31 (7432.49) | 66596.96 (7630.98) | 61726.51 (10133.59) | 61575.99 (9038.66) | 67901.35 (10059.43) |
| L cuneus | 43061.31 (6805.1) | 43813.26 (9743.69) | 43011.29 (8442.26) | 41803.14 (7420.29) | 41019.24 (12460.97) | 44123.91 (4415.42) | 42634.64 (8668.02) | 41456.17 (8625.88) | 40911.49 (6834) | 43835.3 (7348.74) |
| L entorhinal | 22609.84 (6329.62) | 19816.93 (6004.43) | 22021.78 (5225.33) | 20343.35 (5976.53) | 23363.73 (5639.95) | 20263.33 (7034.06) | 21708.47 (5814.26) | 22358.29 (4625.79) | 20758.57 (5947.67) | 19561.63 (4964.37) |
| L frontal pole | 8246.91 (2662.85) | 7651.07 (2551.95) | 6861.36 (2938.93) | 8737.33 (2328.09) | 7654.15 (2977.57) | 10336.71 (3974.95) | 8588.87 (2642.63) | 9034.91 (2857.16) | 7253.21 (2831.14) | 7333.52 (3299.46) |
| L fusiform | 64703.85 (14395.52) | 63592.1 (10392.25) | 64161.01 (10689.48) | 66947.98 (9029.14) | 62336.1 (9489.62) | 68055.27 (8904.78) | 63284.19 (6658.41) | 65994.77 (8028.14) | 63515.85 (9016.67) | 65076.82 (9286.47) |
| L hippocampus | 82284.38 (10817.37) | 75211.38 (9029.25) | 80021.31 (10038.54) | 76582.73 (9296.58) | 82191.26 (11711.06) | 74679.83 (6415.19) | 77496.36 (11562.73) | 77522.28 (9356.27) | 83545.61 (7678.36) | 75499.62 (9252.53) |
| L isthmus cingulate cortex | 40075.55 (6911.84) | 39429.08 (4710.28) | 37933.48 (5029.91) | 40970.89 (4336.9) | 38623.9 (2459.82) | 40026.16 (5368.09) | 39034.66 (3591.14) | 39827.74 (6695.26) | 39430.07 (5253.9) | 39200.32 (4651.64) |
| L insula | 103575.5 (9785.36) | 103711.6 (14635.41) | 102624 (17580.34) | 100756.2 (11552.27) | 103934.6 (7598.45) | 107063.8 (13630.9) | 100282.8 (9884.63) | 105779.8 (14453.28) | 105739.7 (18896.72) | 102481.5 (12473.77) |
| L inferior parietal lobule | 85958.8 (11021.1) | 86302.28 (7293.14) | 86130.68 (11427.54) | 87346.52 (13076.33) | 84582.68 (23653.73) | 91542.85 (7212.73) | 87442.77 (7918.73) | 86018.96 (10649.47) | 82156.05 (11509.91) | 88843.61 (14150.01) |
| L inferior temporal gyrus | 64669.01 (16685.46) | 68228.13 (16276.89) | 65047.99 (12310.55) | 67738.11 (8088.38) | 63932.45 (17760.48) | 71184.53 (10768.38) | 67905.35 (10962.78) | 67516.28 (10763.1) | 62607.82 (10621.66) | 68508.18 (9885.21) |
| L lingual | 56163.35 (10290.29) | 52756.15 (7544.96) | 53753.33 (9384.62) | 56496.7 (8709.53) | 54672.75 (11147.62) | 56413.83 (4854.13) | 54738.79 (8812.12) | 53993.35 (8469.32) | 54592.74 (8958.88) | 52579.94 (8289.36) |
| L lateral orbitofrontal | 64803.14 (11285.9) | 66031.64 (8550.86) | 66167.7 (9319.79) | 63280.91 (7806.77) | 61085.29 (14410.24) | 73404.52 (9030) | 66746.4 (9086.65) | 68270.9 (8408.95) | 64489.8 (9053.79) | 62829.58 (9111.05) |
| L lateral occipital gyrus | 83405.39 (18063.29) | 86630.25 (11695.85) | 83818.6 (9147.94) | 84558.88 (11930.55) | 80170.27 (23079.42) | 91018.84 (8006.8) | 82214.8 (10010.83) | 82750.4 (13464.19) | 77843.31 (14296.17) | 85981.06 (12672.33) |
| L medial orbitofrontal | 55463.02 (10440.02) | 53384.87 (11211.71) | 53511.45 (12217.3) | 53192.8 (9264.37) | 54229.72 (8158.08) | 58990.1 (8316.64) | 57102.36 (12104.75) | 56791.64 (11182.52) | 54699.09 (10765.18) | 50619.68 (10027.84) |
| L middle temporal gyrus | 65222.15 (15220.49) | 63606.16 (8334.49) | 63624.46 (9399.08) | 67447.68 (10525.25) | 65114.49 (16785.54) | 71410.92 (6766.21) | 67050.32 (9063.45) | 66280.68 (10504.75) | 62503.36 (10606.17) | 65061.99 (11320.27) |
| L pallidum | 55291.06 (8017) | 52212.66 (8177.98) | 51165.26 (10251.98) | 55424.29 (9574.5) | 53673.85 (10964.71) | 50371.43 (6983.36) | 50994.23 (7484.76) | 53468.04 (11594) | 57936.89 (9260.4) | 50422.99 (8698.33) |
| L paracentral | 45794.94 (11134.63) | 46674.54 (4013.46) | 45195.18 (6348.97) | 44743.26 (5146.24) | 44096.08 (12375.02) | 46679.33 (4282.45) | 44897.87 (5390.46) | 46476.73 (7130.36) | 47205.75 (4332.71) | 44580.21 (7375.29) |

| STRENGTH | Study 1 | | Study 2 | | Study 3 | | Study 4 | | Study 5 | |
|---|---|---|---|---|---|---|---|---|---|---|
| | HC (N=20) | AN (N=23) | HC (N=25) | AN (N=26) | HC (N=13) | AN (N=16) | HC (N=35) | AN (N=30) | HC (N=26) | AN (N=52) |
| | Mean (SD) | Mean (SD) | Mean (SD) | Mean (SD) | Mean (SD) | Mean (SD) | Mean (SD) | Mean (SD) | Mean (SD) | Mean (SD) |
| L parahippocampal | 34906.31 (5535.98) | 32148.58 (5498.47) | 34546.74 (7133.15) | 32597.78 (3875.34) | 35068.43 (7671.86) | 31095.23 (4214.3) | 33631.75 (5082.4) | 32389.74 (5644.11) | 35298.74 (7876.36) | 31796.98 (5224.72) |
| L posterior cingulate cortex | 37780.12 (10109.98) | 39975.02 (4200.59) | 37704.22 (5290.53) | 38687.02 (5148.39) | 37301.29 (3103.35) | 40534.65 (6038.28) | 38622.31 (5589.51) | 39353.99 (6274.6) | 38607.52 (5553.19) | 38479.56 (6112.22) |
| L precuneus | 80889.21 (15190.91) | 81638.47 (9243.29) | 77408.17 (8189.93) | 81425.33 (8226.08) | 77947.77 (18561.01) | 82327.13 (8522.44) | 78181.18 (6975.36) | 80330.78 (8883.8) | 80580.26 (7745.78) | 78602.48 (9534.73) |
| L pericalcarine | 58009.14 (11091.17) | 59841.53 (14133.56) | 59350.63 (14283.83) | 56158.21 (11532.32) | 55178.1 (17022.86) | 61362.34 (8433.75) | 57842.78 (12347.2) | 56377.72 (12417.13) | 57065.77 (11027.84) | 57479.9 (12460.16) |
| L pars opercularis | 42610.18 (10030.15) | 47875.39 (6727.17) | 44242.49 (7698.35) | 45525.6 (6902.9) | 43336.4 (14854.57) | 49912.86 (7488.49) | 45768.18 (5754.96) | 43295.73 (9042.63) | 43607.78 (9084.38) | 46801.06 (6871.1) |
| L pars orbitalis | 21651.21 (4433.45) | 23062.06 (3365.92) | 22630.46 (4054.76) | 21750.08 (4079.41) | 20221.6 (6005.62) | 25844.96 (3401.58) | 23245.13 (4771.19) | 23107.07 (3735.39) | 20026.12 (3808.63) | 22776.43 (4568.83) |
| L postcentral | 77606.97 (18282.07) | 82563.76 (5401.55) | 75698.95 (10472.06) | 82932.74 (10766.22) | 80177.62 (24161.76) | 86450.31 (9239.2) | 81904.99 (7617.76) | 79323.75 (11084.14) | 80832.67 (9272.88) | 79423.25 (9485.21) |
| L precentral | 119038.7 (25888.21) | 124329.7 (7463.27) | 119034.4 (16356.2) | 122415.6 (14127.26) | 117262 (32461.51) | 130576.6 (8528.62) | 123284.5 (9589.49) | 122736.9 (9114.98) | 121369.3 (8696.49) | 120910.1 (11099.51) |
| L pars triangularis | 37401.83 (6040.36) | 38433.56 (6703.55) | 36976.48 (6140.67) | 38887.72 (5673.67) | 35185.34 (10185.46) | 43359.74 (4715.1) | 40727.5 (5957.47) | 37159.48 (5168.33) | 36204.87 (6498.56) | 38037.8 (6886.68) |
| L putamen | 131347 (10276.41) | 136297.5 (15643.36) | 132219.1 (14418.1) | 132607 (16543.39) | 129723.2 (28256.69) | 137269.3 (16597.15) | 129961.7 (14315.29) | 134501.2 (18090.64) | 135576.5 (15137.05) | 133204.7 (17706.8) |
| L rostral anterior cingulate c ortex | 32979.55 (6252.29) | 32175.93 (5755.79) | 33694.78 (6770.98) | 31737.14 (5847.3) | 32365.01 (3365.58) | 34248.75 (4945.73) | 32634.8 (4430.58) | 34347.55 (8127.06) | 34426.92 (6342.57) | 31010.28 (4936.68) |
| L rostral middle frontal gyru s | 107680.2 (17841.41) | 111583.3 (16481.68) | 114109.3 (25954.95) | 100323.5 (8323.11) | 102127.8 (29041.76) | 121793.7 (8758.29) | 111164.2 (15706.39) | 105607.4 (14597.51) | 101107.4 (14476.3) | 111517.9 (19740) |
| L superior frontal gyrus | 144698.6 (33193.34) | 155218.7 (10527.35) | 151823.3 (18186.76) | 147498.2 (12302.45) | 143899.3 (37061.29) | 160776.8 (12150.05) | 153050 (13816.22) | 151010.8 (15974.6) | 144055.2 (14648.74) | 150631.1 (11351.13) |
| L supramarginal gyrus | 73696.96 (15142.79) | 75395.64 (5803.64) | 74031.23 (8970.66) | 74292.69 (9216.36) | 73926.15 (21614.89) | 79707.83 (8625.42) | 76511.21 (9607.13) | 75676.62 (10335.42) | 74144.59 (9674.12) | 73364.9 (9843.3) |
| L superior parietal lobule | 115542.1 (21111.58) | 121436.4 (14524.11) | 113587.8 (13159.16) | 123308.2 (12760.44) | 116579.7 (32735.11) | 120242.4 (11769.42) | 116928 (10453.3) | 118397.6 (14340.17) | 116139.8 (13255.97) | 120517.3 (15575.93) |
| L superior temporal gyrus | 65662.4 (11905.42) | 64805.71 (7554.12) | 64173.04 (8911.49) | 66455.78 (6989.34) | 64979.03 (15948.42) | 71858.93 (9425.46) | 67024.05 (8973.11) | 67167.06 (8424.89) | 63076.32 (5942.18) | 65801.92 (6490.58) |
| L thalamus | 121032.4 (9926.18) | 121223.7 (9951.05) | 118099.3 (13757.85) | 120513.4 (11151.58) | 120388.8 (11292.24) | 118276.3 (10115.58) | 120411.4 (12305.39) | 118239.7 (16070.86) | 124881.4 (15027.16) | 116261.3 (12476.79) |
| L temporal pole | 14110.13 (5073.2) | 12611.68 (3501.27) | 12994.99 (3385.77) | 13174.09 (3159.53) | 12049.29 (3907.03) | 14937.56 (5960.47) | 13652.63 (3141.94) | 14102.85 (3706.3) | 11838.67 (3845.42) | 12872.89 (3439.32) |
| L transverse temporal | 15289.47 (3110.07) | 15111.44 (2299.12) | 15143.82 (2658.57) | 15351.22 (1903.55) | 14897.97 (4220.61) | 15706.56 (2004.77) | 15712.74 (2181.42) | 15140.29 (2595.01) | 14927.33 (1808.48) | 15525.83 (3293.97) |
| R nucleus accumbens | 20316.21 (4999.94) | 20815.38 (4935.87) | 21073.8 (4024.17) | 20638.11 (5302.43) | 21830.7 (7871.28) | 18653.29 (3757.03) | 20039.51 (5006.94) | 19343.91 (3629.56) | 21732.74 (5475.57) | 22021.83 (4865.72) |
| R amygdala | 32073.07 (6178.5) | 32806.97 (5615.45) | 32988.45 (7412.83) | 28720.42 (3949.93) | 33874.16 (10849) | 31125.5 (5136.87) | 29442.08 (4389.91) | 32726.1 (5235.12) | 33009.78 (4354.83) | 30509.83 (6179.38) |
| R bank of the superior temp oral sulcus | 31337.98 (3896.39) | 33423.32 (6160.59) | 32837.3 (11614.36) | 29874.5 (5155.72) | 33339.58 (7043.91) | 33128.08 (6586.17) | 31308.88 (6586.39) | 32571.52 (9527.46) | 33052.22 (7103.61) | 33012.01 (7068.97) |
| R caudal anterior cingulate | 30497.16 (7606.61) | 32353.96 (4190.1) | 30817.34 (7022.74) | 32359.05 (5178.8) | 32273.43 (7225.84) | 30128.54 (6071.58) | 30675.06 (5540.31) | 31512.08 (5585.36) | 30784.74 (5660.87) | 32291.1 (5550.28) |
| R caudate | 104916.31 (12839.01) | 107447.96 (11762.12) | 104936.15 (17706.04) | 105556.14 (17612.72) | 106641.28 (26235.85) | 99338.74 (7412.13) | 101933.82 (14325.64) | 101958.41 (13026.03) | 113790.74 (16036.44) | 101290.27 (13609.9) |
| R Cerebellum | 48208.22 (11243.12) | 46186.87 (9765.26) | 50447.21 (12227.87) | 47959.91 (9760.5) | 51358.76 (17004.7) | 43317.25 (7337.25) | 51157 (9836.28) | 42821.55 (8325.83) | 48416 (9616.91) | 45098.37 (9357.59) |

| | Study 1 | | Study 2 | | Study 3 | | Study 4 | | Study 5 | |
|---|---|---|---|---|---|---|---|---|---|---|
| STRENGTH | HC (N=20) | AN (N=23) | HC (N=25) | AN (N=26) | HC (N=13) | AN (N=16) | HC (N=35) | AN (N=30) | HC (N=26) | AN (N=52) |
| | Mean (SD) | Mean (SD) | Mean (SD) | Mean (SD) | Mean (SD) | Mean (SD) | Mean (SD) | Mean (SD) | Mean (SD) | Mean (SD) |
| R caudal middle frontal gyrus | 62514.5 (15244.21) | 66330.21 (5813.12) | 65534.47 (11369.96) | 60013.88 (8704.51) | 64165.6 (13876.9) | 65586.74 (10619.93) | 64316.75 (8543.72) | 59563.27 (8466.6) | 60064.07 (9188.09) | 67108.69 (10896.52) |
| R cuneus | 44644.13 (5655.46) | 45816.2 (5328.84) | 46240.01 (10580.42) | 44179.04 (5488.77) | 45033.93 (8007.03) | 46037.47 (8629.32) | 46006.92 (5981.61) | 43726.81 (6418.27) | 43567.69 (6622.52) | 45555.17 (6929.99) |
| R entorhinal | 23650.03 (8477.26) | 20891.42 (5537.83) | 22040.49 (6729.44) | 20182.85 (4290.69) | 25458.13 (9998.38) | 19314.6 (5429.26) | 20335.76 (5683.38) | 22912.31 (4954.62) | 22028.39 (5891.78) | 20106.87 (6855.04) |
| R frontal pole | 10228.79 (4104.63) | 9769.41 (2707.26) | 8465.76 (2936.36) | 11602.61 (2912.32) | 9577.63 (3179.76) | 13188.54 (4695.06) | 10625.19 (3498.9) | 11121.86 (2817.82) | 9890.65 (3885.29) | 9188.18 (3813.85) |
| R fusiform | 64544.8 (12439.37) | 67236.92 (7725.27) | 67282.14 (10923.88) | 65780.3 (7870.86) | 67840.37 (16172.7) | 65186.76 (7662.39) | 62809.52 (6214.37) | 67257.13 (7915.62) | 65170.08 (9874.34) | 67957.57 (10972.08) |
| R hippocampus | 90171.46 (15841.84) | 83929.66 (14824.28) | 87338.84 (10560.23) | 81307.55 (9358.92) | 89590.66 (19382.9) | 79911.09 (4612.33) | 83464.95 (10052.34) | 83193.5 (11765.68) | 88517.67 (8242.04) | 83345.39 (12874.99) |
| R isthmus cingulate cortex | 36979.77 (6682.58) | 36352.66 (2326.56) | 34986.54 (4744.19) | 37748.26 (3633.51) | 37188.14 (8135.77) | 37204.22 (5013.91) | 35534.55 (4079.89) | 36849.6 (6084.34) | 35942.92 (4861.45) | 36281.75 (4408.77) |
| R insula | 110118.3 (15762.82) | 107014.6 (16839.67) | 103962.7 (24330.04) | 102058.5 (12898.33) | 118866 (24513.07) | 102287.8 (16417.87) | 104474.6 (12309.99) | 108275.9 (18373.47) | 104900.4 (18641.67) | 106774.9 (18502.15) |
| R inferior parietal lobule | 91957.77 (11586.48) | 94105.36 (9682.99) | 93119.51 (11584.76) | 91298.83 (10692.62) | 98034.31 (14873.51) | 94379.65 (10788.22) | 92004.23 (10528.36) | 95417.51 (10306.24) | 92582.48 (12150.58) | 94517.61 (9494.8) |
| R inferior temporal gyrus | 63678.54 (16159.49) | 68723.34 (10978.53) | 67220.07 (15288.14) | 64325.7 (6263.81) | 70475.22 (15226.99) | 67112.68 (7434.99) | 65579.75 (9279.37) | 67608.38 (11958.98) | 63293.85 (9278.56) | 65982.52 (10307.79) |
| R lingual | 61349.4 (8694.72) | 56634.56 (6424.17) | 59154.12 (10463.46) | 60295.68 (7728.79) | 62053.42 (13466.09) | 57310.12 (4388.04) | 58072.36 (8022.54) | 58669.71 (6636.98) | 58293.65 (6634.73) | 58215.92 (8191.01) |
| R lateral orbitofrontal | 69862.28 (10374.55) | 74153.19 (9407.53) | 73921.57 (9220.4) | 68220.57 (9137.55) | 66303.16 (11561.36) | 77449.9 (10520.16) | 69334.82 (10241.02) | 71872.04 (13941.12) | 74108.83 (13870.09) | 68812.84 (10550.21) |
| R lateral occipital gyrus | 84610.56 (9021.44) | 89253.56 (13804.2) | 88159.6 (8956.39) | 84768.9 (6069.17) | 86367.34 (15650.4) | 90217.86 (7240.55) | 83588.46 (11548.08) | 87420.33 (10223.52) | 80222.42 (7508.28) | 91496.5 (10808.39) |
| R medial orbitofrontal | 46958.67 (8767.99) | 49115.85 (9895.68) | 47732.81 (6922.24) | 47162.36 (7484.3) | 45958.14 (10177.82) | 53035.72 (8353.98) | 49830.68 (8469.6) | 49791.14 (7986.4) | 48907.17 (9830.14) | 45518.13 (8878.16) |
| R middle temporal gyrus | 66556.04 (17770.97) | 72465.88 (13663.17) | 69823.66 (12404.09) | 68997.4 (7410.42) | 72780.93 (12802.85) | 74195.81 (6626.5) | 71097.7 (9618.2) | 70997.14 (8740.27) | 69471.92 (10223.63) | 68549.05 (11350.53) |
| R pallidum | 55205.06 (8614.69) | 56660.77 (8699.98) | 51491.78 (10935.24) | 57966.94 (9459.91) | 56451.47 (18577.13) | 51076.67 (7129.15) | 51427.2 (6838.13) | 55397.07 (12375.92) | 58486.86 (10044.77) | 53088.5 (9348.14) |
| R paracentral | 50367.89 (12304.99) | 49867.07 (3909.79) | 48551.32 (4750.98) | 49745.99 (5796.77) | 51327.75 (9789.11) | 49392.45 (3859.25) | 49539.52 (5465.58) | 49864.55 (6853.84) | 51197.02 (6378.38) | 48950 (6302.67) |
| R parahippocampal | 36638.67 (6472.4) | 33612.77 (6337.06) | 35066.08 (6340.46) | 33091.43 (5664.92) | 37226.72 (9769.26) | 32855.96 (5576.17) | 34285.81 (5414.79) | 33836.93 (4707.33) | 36457.95 (6443.85) | 32588.03 (6110.77) |
| R posterior cingulate cortex | 38734.92 (9093.02) | 39339.22 (4324.85) | 38914.72 (5203.13) | 39039.38 (5123.38) | 40095.88 (7359.7) | 38882.99 (7629.01) | 38032.89 (5367.26) | 40597.15 (6787.28) | 38963.99 (5651.26) | 39368.2 (5545.91) |
| R precuneus | 82691.66 (14950.69) | 82844.71 (8143.21) | 80335.19 (7638.04) | 83211.42 (8850.75) | 83662.69 (13047.63) | 80638.79 (7612.04) | 79084.89 (7855.21) | 80393.51 (7088.11) | 82285.72 (7376.33) | 82748.96 (10399.74) |
| R pericalcarine | 62510.96 (10935.25) | 61667.31 (10978.7) | 62979.61 (11407.35) | 61178.92 (11225.04) | 62751.87 (13875.2) | 61905.31 (10216.92) | 60421.95 (10458.98) | 60906.07 (10513.61) | 61639.32 (8762.23) | 61862.91 (12088.84) |
| R pars opercularis | 43322.08 (11677.33) | 44751.19 (6035.77) | 43005.68 (4970.07) | 42354.42 (6914.21) | 45112.66 (8094.82) | 45276.2 (8839.98) | 42058.07 (5719.79) | 41275.79 (8271.59) | 42603.74 (6596.39) | 45031.32 (7167.2) |
| R pars orbitalis | 25472.94 (5944.76) | 26501.35 (5015.06) | 26298.34 (4612.07) | 25143.85 (4524.7) | 25737.38 (6104.5) | 27595.84 (6563.24) | 27156.83 (5383.33) | 25580.09 (4312.41) | 24457.59 (4619.5) | 26627.89 (4576.24) |
| R postcentral | 79245.57 (17826.97) | 83463.96 (6674.14) | 79073.64 (9594.87) | 81413.38 (7285.62) | 85385.96 (14446.48) | 82318.13 (15446.64) | 78918.97 (9170.59) | 79620.92 (8878.63) | 84040.35 (10907.01) | 81339.78 (9423.23) |
| R precentral | 120900.4 (26470.09) | 124371.3 (7405.57) | 119223.4 (15423.39) | 121980.6 (8419.05) | 124660.6 (17654.95) | 124925.6 (14606.7) | 120692.9 (8760.82) | 120351.3 (11750.74) | 122005.6 (10628.87) | 122993.2 (10844.04) |

| | | | | | | | | | | |
|---|---|---|---|---|---|---|---|---|---|---|
| R pars triangularis | 40341.68 (6323.82) | 41557.84 (7319.26) | 39780.45 (5837.27) | 40568.3 (8038.63) | 42722.85 (10124.97) | 41675.83 (8454.49) | 41338.6 (5454.61) | 39757.89 (7401.62) | 38631.37 (4884.81) | 41471.52 (7550.08) |
| R putamen | 132270.4 (13080.37) | 148974.4 (24381.13) | 136997 (17206.72) | 135806 (17764.22) | 143383.8 (29329.98) | 131781.9 (19676.62) | 132882.5 (15931.93) | 136604.4 (14824.93) | 138032.6 (14599.1) | 142814.1 (18364.09) |
| R rostral anterior cingulate c ortex | 26321.1 (3020.3) | 27646.33 (3380.07) | 27446.97 (6244.73) | 27566.48 (4829.77) | 28069.5 (6959.73) | 26540.68 (5810.32) | 27370.13 (3989.95) | 27940.21 (4781.96) | 27140.15 (5474.38) | 27283.5 (4722.13) |
| R rostral middle frontal gyru s | 112793.6 (19472.04) | 117693.1 (18924.88) | 120132.2 (27301.42) | 105359 (11692.27) | 115242.2 (23440.22) | 119856.3 (9459) | 113049.8 (14371.85) | 110653.1 (14209.19) | 107703.8 (18519.38) | 116140.6 (20477.54) |
| R superior frontal gyrus | 141236.8 (31876.8) | 151003.6 (11905.33) | 145452.8 (14847) | 146633.1 (14461.24) | 145480.2 (25132.64) | 151320.1 (13315.16) | 147668.9 (12334.23) | 143923.8 (14465.04) | 139965.7 (13452.32) | 147624.6 (13298.65) |
| R supramarginal gyrus | 71075.8 (15687.57) | 73351.27 (6400.65) | 70223.66 (10667.83) | 72385.38 (7925.74) | 74912.24 (12565.34) | 75082.33 (10688.27) | 72257.19 (9074.35) | 72130.21 (9577.23) | 71349.32 (8623.04) | 71448.98 (11559.23) |
| R superior parietal lobule | 111811.9 (26217.03) | 119517.2 (9160.8) | 111083.2 (11531.47) | 118089.5 (11594.01) | 120502.2 (17919.24) | 113379.5 (14470.94) | 113894.4 (8673.85) | 116886.8 (15430.95) | 112695.9 (13110.52) | 120106.7 (13859.62) |
| R superior temporal gyrus | 60028.06 (11272.3) | 63202.83 (11372.98) | 59826.42 (8340.94) | 60175.24 (6823.11) | 61297.63 (10538.84) | 64851.81 (12361.9) | 61455.22 (7019.52) | 60721.04 (9188) | 60265.25 (8481.18) | 60115.82 (9663.38) |
| R thalamus | 113711.4 (9833.97) | 110136.6 (10304.26) | 110215.2 (11065.48) | 110303.3 (10242.2) | 115707 (20960.67) | 106884.7 (8419.93) | 111798.6 (11372.04) | 109426.6 (11118.28) | 115921.5 (11144.51) | 105693.5 (10802.8) |
| R temporal pole | 15146.91 (7102.57) | 14882.51 (3954.27) | 14100.83 (3441.24) | 14035.56 (2646.77) | 16737.57 (7715.26) | 12887.53 (5444.28) | 14984.15 (3841.16) | 14810.35 (4124.81) | 12407.18 (4100.37) | 14838.06 (3901.48) |
| R transverse temporal | 13514.33 (2767.58) | 13556.71 (2178.87) | 13383.98 (2539.32) | 13192.02 (2071.72) | 13702.36 (2811.76) | 13758.5 (2173.5) | 14035.48 (2082.31) | 13353.35 (2467.76) | 13386.72 (2675.71) | 13095.86 (2175.24) |
| EFFICIENCY | **Study 1** | | **Study 2** | | **Study 3** | | **Study 4** | | **Study 5** | |
| | HC (N=20) | AN (N=23) | HC (N=25) | AN (N=26) | HC (N=13) | AN (N=16) | HC (N=35) | AN (N=30) | HC (N=26) | AN (N=52) |
| | Mean (SD) | Mean (SD) | Mean (SD) | Mean (SD) | Mean (SD) | Mean (SD) | Mean (SD) | Mean (SD) | Mean (SD) | Mean (SD) |
| L nucleus accumbens | 106.64 (25.59) | 109.78 (26.28) | 107.85 (21.08) | 108.05 (29.28) | 105.03 (25.8) | 105.67 (23.81) | 105.24 (23) | 98.96 (23.7) | 115.96 (25.06) | 112.92 (26.89) |
| L amygdala | 131.27 (15.21) | 139.02 (21.16) | 139.03 (24.87) | 129.05 (19.46) | 140.66 (19.23) | 137.65 (26.77) | 135.01 (26.29) | 137.73 (20.93) | 143.8 (30.01) | 130.14 (20.9) |
| L bank of the superior temp oral sulcus | 125.78 (23.89) | 142.28 (24.81) | 134.78 (28.03) | 134.21 (13.17) | 129.98 (34.33) | 145.46 (25.38) | 132.43 (31.97) | 140.34 (29.63) | 138.74 (25.35) | 137.97 (33.8) |
| L caudal anterior cingulate | 181.97 (48.76) | 192.89 (36.29) | 182.61 (26.9) | 191.01 (25.66) | 182.6 (21.84) | 197.18 (37.98) | 181.85 (27.67) | 192.25 (35.7) | 184.73 (32.29) | 183.69 (30.2) |
| L caudate | 539.69 (103.31) | 546.6 (73.62) | 537.69 (77.84) | 547.33 (97.17) | 515.38 (119.46) | 555.66 (63.36) | 531.35 (81.49) | 526.63 (68.81) | 579.91 (91.78) | 526.48 (77.3) |
| L Cerebellum | 182.46 (28.6) | 176.08 (34.72) | 187.35 (46.2) | 185.15 (42.64) | 171.99 (36.78) | 201.72 (46.7) | 195.46 (31.66) | 174.94 (38.18) | 179.79 (46.91) | 172.08 (46.69) |
| L caudal middle frontal gyru s | 324.87 (81.86) | 351.74 (39.82) | 335.29 (52.13) | 322.31 (47.27) | 325.81 (95) | 356.77 (39.57) | 354.2 (43.76) | 324.86 (37.64) | 326.72 (48.2) | 330.8 (47.97) |
| L cuneus | 120.7 (18.67) | 122.77 (40.82) | 124.65 (34.16) | 114.58 (19.08) | 110.31 (36.79) | 134.6 (24.13) | 126.83 (25.11) | 117.86 (25.24) | 113.08 (22.51) | 125.73 (25.13) |
| L entorhinal | 70.09 (21.58) | 59.53 (16.13) | 69.57 (20.97) | 60.16 (15.89) | 70.22 (21.42) | 65.68 (23.6) | 68.34 (15.63) | 68.57 (16.51) | 65.47 (23.49) | 59.02 (14.93) |
| L frontal pole | 40.3 (11.22) | 37.66 (12.29) | 33.39 (15.65) | 43.37 (12.11) | 39.96 (16.58) | 49.07 (22.77) | 43.11 (11.25) | 43.24 (11.84) | 34.38 (11.84) | 36.79 (16.76) |
| L fusiform | 245.97 (38.89) | 262.1 (52.84) | 258.7 (49.58) | 261.7 (37.47) | 241.62 (42.55) | 270.23 (44.71) | 256.35 (39) | 256.04 (49.23) | 255.16 (53.71) | 256.49 (45.03) |
| L hippocampus | 382.73 (40.03) | 356.48 (44.62) | 372.46 (48.02) | 366.94 (55.44) | 365.63 (35.8) | 373.86 (37.09) | 386.23 (51.36) | 356.74 (54.97) | 377.69 (46.61) | 358.09 (52.38) |
| L isthmus cingulate cortex | 303.9 (55.18) | 312.29 (42.76) | 293.67 (52.78) | 316.62 (40.46) | 303.13 (28.37) | 312.14 (41.57) | 308.69 (32.04) | 304.33 (59.88) | 304.76 (55.49) | 299.93 (42.52) |
| L insula | 604.52 (75.13) | 634.54 (82.88) | 608.42 (94.84) | 602.9 (82.25) | 611.87 (58.85) | 632.43 (80.27) | 601.92 (64) | 619.76 (88.48) | 616.51 (105.79) | 610.59 (67.39) |
| L inferior parietal lobule | 425.49 (70.31) | 442.05 (41.63) | 423.23 (58.5) | 453 (63.22) | 427.67 (117.6) | 462.75 (66.62) | 450.21 (60.36) | 434.78 (58.99) | 429.5 (59.82) | 442.1 (62.54) |
| L inferior temporal gyrus | 302.83 (57.85) | 333.96 (53.14) | 306.52 (49.46) | 323.87 (43.12) | 308.91 (84.41) | 338.48 (54.71) | 321.18 (55.35) | 327.02 (65.85) | 312.31 (64.94) | 321.16 (46.42) |

| EFFICIENCY | Study 1 | | Study 2 | | Study 3 | | Study 4 | | Study 5 | |
|---|---|---|---|---|---|---|---|---|---|---|
| | HC (N=20) | AN (N=23) | HC (N=25) | AN (N=26) | HC (N=13) | | HC (N=20) | AN (N=24) | HC (N=25) | AN (N=26) |
| | Mean (SD) | Mean (SD) | Mean (SD) | Mean (SD) | Mean (SD) | | Mean (SD) | Mean (SD) | Mean (SD) | Mean (SD) |
| L lingual | 185.4 (25.84) | 186.99 (32.24) | 191.58 (38.49) | 190.71 (29.67) | 184.8 (33.67) | 195.05 (27.54) | 192.97 (30.91) | 183 (36.76) | 185.75 (28.58) | 179.75 (27.93) |
| L lateral orbitofrontal | 260.71 (31.69) | 280.48 (34.47) | 268.64 (42.41) | 273.44 (37.4) | 263.35 (55.01) | 302.57 (42.37) | 275.25 (43.41) | 289.7 (38.89) | 256.19 (44.62) | 268.56 (35.48) |
| L lateral occipital gyrus | 316.65 (46.71) | 308.08 (40.9) | 315.28 (43.69) | 313.78 (48.43) | 300.94 (97.59) | 352.6 (43.4) | 321.63 (44.15) | 322.77 (72.22) | 300.17 (59.17) | 312.77 (60) |
| L medial orbitofrontal | 301.25 (68.31) | 276.78 (84.48) | 267.77 (85.35) | 295.76 (77.92) | 298.39 (42.81) | 307.67 (61.03) | 311.81 (84.92) | 293.84 (75.68) | 278.07 (74.37) | 271.18 (79.9) |
| L middle temporal gyrus | 259.67 (51.18) | 268.12 (39.92) | 255.88 (51.44) | 288.47 (37.64) | 276 (70.82) | 285.05 (57.11) | 273.14 (40.23) | 281.26 (53.18) | 260.18 (41.12) | 257.35 (54.58) |
| L pallidum | 414.21 (75.84) | 413.47 (61.05) | 391.51 (90.55) | 432.62 (71.29) | 414.29 (84.53) | 403.92 (48.92) | 412.65 (54.43) | 420.53 (83.66) | 427.24 (49.25) | 392.86 (67.05) |
| L paracentral | 232.38 (57.75) | 246.28 (19.36) | 233.59 (34.85) | 231.62 (29.71) | 235.42 (66.93) | 248.88 (24.5) | 237.96 (28.81) | 239.89 (31.22) | 236.09 (25.59) | 227.87 (37.96) |
| L parahippocampal | 145.94 (36.02) | 137.23 (29.25) | 145.27 (38.66) | 142.14 (32.55) | 141.2 (31.35) | 143.27 (33.64) | 151.03 (33.4) | 138.62 (35.38) | 144.42 (37.98) | 133.78 (31.64) |
| L posterior cingulate cortex | 271.78 (76.81) | 293.66 (37.33) | 275.83 (55.41) | 275.98 (35.69) | 275.45 (32.33) | 289.09 (37.77) | 282.32 (42.86) | 273.19 (48.71) | 273.32 (44.41) | 274.5 (45.95) |
| L precuneus | 486.74 (103.04) | 514.52 (53.01) | 486.05 (55.95) | 493.88 (44.2) | 482.7 (111.4) | 518.81 (59.29) | 506.66 (46.21) | 493.34 (76.2) | 492.62 (61.75) | 478.54 (57.41) |
| L pericalcarine | 141.98 (27.31) | 151.17 (40.74) | 152.82 (36.26) | 140.92 (28.49) | 131.38 (43.71) | 163.5 (17.9) | 153.59 (29.61) | 142.99 (41.73) | 143.77 (22.51) | 145.09 (31.58) |
| L pars opercularis | 241.95 (64.44) | 268.08 (28.69) | 243.96 (48.82) | 259.49 (32.4) | 236.71 (68.49) | 280.23 (26.63) | 262.3 (34.01) | 255.46 (53.62) | 241.15 (49.82) | 252.03 (37.32) |
| L pars orbitalis | 86.96 (20.26) | 91.23 (18.41) | 85.45 (20.85) | 90.97 (17.84) | 83.93 (27.77) | 102.61 (17.37) | 92.81 (17.67) | 95.85 (17.36) | 76.9 (19.94) | 92.52 (20.73) |
| L postcentral | 309.49 (74.96) | 339.68 (26.44) | 310.78 (58.4) | 335.57 (42.7) | 311.24 (86.42) | 351.16 (29.72) | 336.74 (29.5) | 322.02 (49.9) | 315.48 (45) | 319.97 (47.55) |
| L precentral | 539.82 (124.5) | 564.12 (43.64) | 540.26 (92.04) | 546.55 (61.02) | 521.09 (137.2) | 586.57 (50.6) | 563.89 (48.19) | 551.75 (70.08) | 537.89 (60.44) | 534.97 (59.81) |
| L pars triangularis | 174.7 (41.7) | 183.25 (23.06) | 168.8 (33.77) | 187.97 (26.13) | 174.6 (48.83) | 195.99 (26.38) | 191.39 (29.39) | 186.33 (34.24) | 168.14 (28.45) | 177.98 (29.52) |
| L putamen | 746.89 (82.47) | 787.47 (63.83) | 753.69 (84.86) | 778.1 (93.48) | 747 (160) | 806.6 (85.82) | 772.09 (74.39) | 787.74 (91.02) | 758.53 (78.71) | 766.57 (97.26) |
| L rostral anterior cingulate cortex | 197.31 (59.53) | 184.77 (29.91) | 191.78 (43.22) | 183.13 (41.83) | 194.96 (39.07) | 201.82 (36.77) | 193.03 (34.7) | 200.21 (53.65) | 190.23 (54.15) | 182.84 (38.46) |
| L rostral middle frontal gyrus | 498.45 (104.9) | 511.3 (53.13) | 491.9 (71.55) | 491.16 (58.37) | 475.66 (133.15) | 562 (72.3) | 513.68 (58.35) | 511.64 (68.99) | 484.46 (55.66) | 496.96 (53.01) |
| L superior frontal gyrus | 728.65 (163.91) | 770.8 (61.98) | 742.01 (98.36) | 755.12 (69.95) | 733.51 (170.99) | 791.89 (80.71) | 768.59 (75.12) | 764.78 (92.59) | 732.43 (72.19) | 737.42 (79.69) |
| L supramarginal gyrus | 361.23 (77.02) | 376.33 (25.78) | 367.23 (52.82) | 368.6 (53.8) | 361.69 (101.36) | 394.72 (45.83) | 380.4 (43.1) | 371.35 (59.62) | 365.56 (59.12) | 357.89 (56.33) |
| L superior parietal lobule | 573.07 (110.52) | 614.22 (60.06) | 579.91 (75.29) | 625.13 (63.88) | 582.7 (158.86) | 626.93 (71.42) | 619.3 (53.57) | 611.54 (93.59) | 585.66 (63.56) | 601.49 (67.06) |
| L superior temporal gyrus | 310.13 (38.97) | 324.76 (26.22) | 310.57 (61.25) | 328.19 (33.65) | 308.29 (79.25) | 354.04 (42.47) | 334.01 (42.53) | 322.86 (47.74) | 312.45 (36.78) | 315.69 (47.1) |
| L thalamus | 899.56 (109.61) | 911.76 (77.21) | 892.8 (100.25) | 901 (77.13) | 895.29 (77.53) | 912.85 (74.04) | 911.11 (85.53) | 894.54 (128.39) | 914.44 (98.9) | 884.77 (85.44) |
| L temporal pole | 51.31 (18.44) | 44.14 (10.8) | 42.64 (14.9) | 44.08 (10.53) | 43.79 (11.71) | 50.6 (15.8) | 47.31 (11.06) | 51.04 (14.6) | 42.94 (12.96) | 44.25 (12.08) |
| L transverse temporal | 80.71 (15.72) | 84.96 (9.41) | 80.69 (17.03) | 85.7 (14.74) | 81.35 (24.76) | 86.9 (10.35) | 88.79 (13.7) | 83.63 (17.46) | 86.84 (20.45) | 84.31 (16.99) |
| R nucleus accumbens | 123.88 (24.79) | 125.2 (37.71) | 126.26 (23.63) | 128.03 (41.64) | 134.34 (37.9) | 115.82 (32.13) | 123.25 (33.76) | 117.22 (33.17) | 131.49 (29.5) | 129.21 (31.22) |
| R amygdala | 153.01 (29.47) | 165.82 (27.72) | 161.76 (29.78) | 145.3 (20.22) | 165.44 (35.43) | 160.66 (28.69) | 153.36 (20.68) | 155.99 (28.88) | 163.64 (21.44) | 149.69 (30.85) |

| EFFICIENCY | Study 1 | | Study 2 | | Study 3 | | Study 4 | | Study 5 | |
|---|---|---|---|---|---|---|---|---|---|---|
| | HC (N=20) | AN (N=23) | HC (N=25) | AN (N=26) | HC (N=13) | AN (N=16) | HC (N=35) | AN (N=30) | HC (N=26) | AN (N=52) |
| | Mean (SD) | Mean (SD) | Mean (SD) | Mean (SD) | Mean (SD) | Mean (SD) | Mean (SD) | Mean (SD) | Mean (SD) | Mean (SD) |
| R bank of the superior temp oral sulcus | 138.33 (25.73) | 159.88 (31.4) | 142.22 (36.84) | 147.36 (28.94) | 150.57 (24.68) | 151.53 (23.16) | 147.05 (35.27) | 146.95 (34.82) | 149.11 (27.54) | 154.76 (31.72) |
| R caudal anterior cingulate | 206.79 (51.23) | 213.71 (32.93) | 197.99 (44.25) | 214.15 (26.08) | 204.5 (28.26) | 207.79 (41.1) | 207.35 (32.24) | 209.96 (39.24) | 202.91 (30.77) | 209.8 (32.79) |
| R caudate | 583.25 (90.53) | 594.94 (74.2) | 583.76 (106.22) | 591.58 (112.93) | 587.26 (115.55) | 567.95 (61.6) | 577.99 (82.82) | 571.24 (81.86) | 628.63 (89.66) | 566.16 (78.47) |
| R Cerebellum | 179 (33.34) | 173.94 (38.71) | 182.96 (52.95) | 184.5 (39.75) | 172.11 (54.17) | 185.07 (36.88) | 193.33 (39.64) | 170.25 (35.19) | 172.99 (43.78) | 170.35 (42.51) |
| R caudal middle frontal gyru s | 319.65 (79.63) | 349.97 (38.05) | 341.22 (52.11) | 311.49 (43.59) | 337.83 (43.2) | 344.54 (43.47) | 332.1 (41.26) | 318.63 (45.16) | 316.24 (39.54) | 335.53 (47.43) |
| R cuneus | 122.95 (16.39) | 123.97 (25.54) | 126.9 (29.49) | 123.57 (18.22) | 119.52 (23.74) | 133.99 (29) | 131.53 (21.67) | 120.17 (24.19) | 119.71 (22.63) | 123.32 (22.26) |
| R entorhinal | 68.03 (28.26) | 58.01 (22.68) | 63.88 (17.4) | 60.08 (18.79) | 67.82 (26.25) | 66.38 (22.56) | 64.89 (20.36) | 65.2 (16.69) | 59.31 (19.7) | 57.97 (18.87) |
| R frontal pole | 44.55 (11.86) | 44.27 (13.93) | 37.11 (15.71) | 53.81 (14.34) | 44.55 (10.33) | 56.09 (21.07) | 48.77 (14.89) | 51.42 (17.28) | 42.79 (14.6) | 41.81 (17.84) |
| R fusiform | 225.93 (33.35) | 254.44 (47.68) | 248.01 (49.49) | 229.99 (33.14) | 245.12 (53.13) | 236.69 (31.82) | 232.74 (34.79) | 241.69 (46.06) | 242.57 (42.33) | 251.65 (46.41) |
| R hippocampus | 437.81 (60.01) | 435.49 (76.12) | 439.83 (50.04) | 418.93 (41.66) | 437.11 (56.87) | 421.94 (30.69) | 440.6 (61.39) | 413.87 (67) | 433.31 (47.11) | 424.29 (58.94) |
| R isthmus cingulate cortex | 266.53 (57.5) | 271.73 (28.51) | 259.61 (49.18) | 279.38 (32.04) | 279.08 (49.72) | 283.51 (46.04) | 270.02 (38.22) | 273.61 (47.9) | 263.91 (48.77) | 265.35 (40.1) |
| R insula | 646.87 (83.44) | 621.32 (73.32) | 607.13 (113.76) | 609.88 (100.36) | 701.31 (112.42) | 602.67 (84.01) | 632.42 (75.07) | 624.67 (103.59) | 614.97 (118.74) | 616.65 (89.38) |
| R inferior parietal lobule | 474.51 (91.1) | 506.08 (56.68) | 477.96 (63.94) | 485.44 (67.02) | 501.52 (65.95) | 489.22 (68.4) | 485.11 (54.08) | 502.43 (71.82) | 493.77 (64.2) | 486.08 (62.28) |
| R inferior temporal gyrus | 279.42 (36.61) | 307.4 (42.3) | 286.12 (48.56) | 297.83 (38.33) | 305.58 (44.62) | 298.33 (51.19) | 295.04 (40.06) | 295.56 (63.21) | 289.72 (49.17) | 283.86 (42.33) |
| R lingual | 209.81 (24.28) | 189.26 (28.1) | 202.93 (44.13) | 204.96 (28.09) | 202.15 (42.45) | 203.88 (20.05) | 203.12 (33.91) | 198.16 (34.25) | 199.97 (23.74) | 195.84 (37.62) |
| R lateral orbitofrontal | 295.54 (66.73) | 343.41 (57.16) | 321.98 (68.41) | 314.91 (65.15) | 288.71 (51.88) | 356.82 (75.06) | 311.53 (59.69) | 321.81 (68.32) | 318.34 (77.42) | 314.14 (62.14) |
| R lateral occipital gyrus | 302.57 (41.04) | 338.55 (46.1) | 321.72 (54.28) | 310.25 (34.81) | 322.7 (44.65) | 334.85 (52.4) | 324.32 (51.48) | 330.36 (64.92) | 301.65 (49.68) | 323.6 (48.88) |
| R medial orbitofrontal | 211.29 (61.48) | 227.07 (66.41) | 211.32 (53.45) | 212.85 (51.03) | 205.84 (31.76) | 243.26 (52.22) | 234.09 (73.53) | 221.17 (46.38) | 223.55 (69.03) | 212.35 (58.57) |
| R middle temporal gyrus | 277.8 (53.34) | 292.29 (37.25) | 276.8 (63.88) | 309.45 (28.26) | 289.24 (52.07) | 306.12 (54.4) | 298.81 (31.77) | 297.39 (57.89) | 280.9 (43.62) | 274.81 (58.45) |
| R pallidum | 407.31 (66.66) | 422.68 (49.32) | 385.25 (88.43) | 436.47 (65.41) | 417.28 (103.53) | 407.03 (49.86) | 407.82 (49.63) | 420.71 (81.8) | 427.26 (57.35) | 401.42 (62.82) |
| R paracentral | 261.41 (66.71) | 273.93 (27.31) | 261.27 (30.92) | 267.1 (39.44) | 269.59 (27.53) | 267.61 (20.23) | 268.85 (29.24) | 265.77 (44.36) | 263.98 (28.38) | 256.48 (39.14) |
| R parahippocampal | 156.34 (37.7) | 154.4 (37.1) | 154.41 (33.45) | 150.85 (38.19) | 169.61 (48.15) | 149.69 (44.29) | 160.28 (34.67) | 149.61 (37.66) | 159.62 (40.5) | 140.17 (30.22) |
| R posterior cingulate cortex | 279.59 (67.77) | 285.1 (38.1) | 283.13 (42.17) | 280.53 (39.36) | 287.14 (54.16) | 281.48 (52.29) | 268.91 (43.06) | 285.62 (46.82) | 280.17 (46.47) | 280.19 (42.43) |
| R precuneus | 492.14 (96.5) | 497.77 (44.1) | 487.88 (61.01) | 498.53 (45.73) | 495.75 (38.18) | 510.66 (47.49) | 496.02 (50.49) | 486.68 (57.84) | 487.49 (54.27) | 484.21 (50.62) |
| R pericalcarine | 146.41 (23.8) | 141.55 (23.14) | 149.75 (31) | 146.43 (26.18) | 140.47 (26.6) | 158.01 (22.52) | 150.18 (29.97) | 146.74 (35.4) | 144.83 (20.25) | 144.23 (30.39) |
| R pars opercularis | 230.55 (64.08) | 238.56 (30.25) | 223.25 (41.41) | 233.76 (30.75) | 236.79 (23.79) | 231.86 (24.83) | 230.55 (36.12) | 233.56 (47.59) | 229.78 (36.67) | 234.88 (40.59) |
| R pars orbitalis | 98.2 (23.43) | 101.63 (29.48) | 98.6 (24.59) | 104.41 (23.63) | 102.13 (23.45) | 108.2 (30.7) | 108.81 (21.79) | 105.38 (21.49) | 88.02 (17.47) | 104.68 (24.18) |
| R postcentral | 316.7 (72.84) | 339.71 (41.69) | 322.84 (40.48) | 332.37 (37.05) | 335.75 (36.17) | 323.1 (26.75) | 328.77 (35.35) | 324.45 (52.37) | 337.68 (47.27) | 317.79 (48.33) |

| EFFICIENCY | Study 1 | | Study 2 | | Study 3 | | Study 4 | | Study 5 | |
|---|---|---|---|---|---|---|---|---|---|---|
| | HC (N=20) | AN (N=23) | HC (N=25) | AN (N=26) | HC (N=13) | AN (N=16) | HC (N=35) | AN (N=30) | HC (N=26) | AN (N=52) |
| | Mean (SD) | Mean (SD) | Mean (SD) | Mean (SD) | Mean (SD) | | Mean (SD) | Mean (SD) | Mean (SD) | Mean (SD) |
| R precentral | 537.19 (122.53) | 561.94 (42.16) | 542.91 (64.55) | 544.06 (45.98) | 548.86 (59.82) | 562.37 (31.69) | 555.35 (45.57) | 549.48 (75.98) | 542.05 (57.42) | 538.42 (67.86) |
| R pars triangularis | 188.31 (43.11) | 193.88 (40.18) | 182.8 (36.31) | 198.8 (29.37) | 199.39 (32.3) | 191.15 (32.9) | 196.93 (36.64) | 200.4 (40.59) | 182.16 (31.44) | 194.05 (38) |
| R putamen | 756.24 (101.09) | 837.36 (83.16) | 772.21 (103.67) | 809.89 (94.28) | 802.99 (126.47) | 789.57 (103.49) | 795.47 (88.57) | 788.49 (105.71) | 776.25 (84) | 809.75 (98.06) |
| R rostral anterior cingulate c ortex | 147.16 (21.48) | 147.55 (17.88) | 141.58 (23.9) | 148.6 (21.19) | 153.77 (28.32) | 151.01 (29.67) | 152.73 (21.71) | 150.36 (23.68) | 142.85 (24.32) | 150.22 (28.23) |
| R rostral middle frontal gyru s | 510.35 (112.53) | 541.32 (83.22) | 521.31 (67.57) | 510.33 (48.58) | 527.66 (68.7) | 556.25 (59.55) | 523.68 (63.8) | 539.02 (80.67) | 515.39 (75.82) | 518.81 (67.12) |
| R superior frontal gyrus | 719.66 (159.31) | 762.43 (67.09) | 723.07 (100.44) | 754.53 (73.94) | 730.98 (86.27) | 766.54 (82.71) | 751.94 (73.78) | 751.52 (81.75) | 728.22 (73.82) | 728.15 (86.92) |
| R supramarginal gyrus | 352.95 (81.93) | 360.11 (30.68) | 349.13 (45.89) | 360.01 (40.83) | 370.46 (55.36) | 360.65 (48.92) | 367.19 (45.94) | 353.85 (55.52) | 349.92 (43.75) | 347.31 (68.76) |
| R superior parietal lobule | 579.03 (140.14) | 617.7 (55.19) | 565.41 (59.41) | 615.76 (63.31) | 611.77 (64.7) | 606.62 (62.17) | 616.88 (48.14) | 613.57 (87.6) | 589.98 (58.55) | 599.86 (58.7) |
| R superior temporal gyrus | 295.67 (49.88) | 312.71 (41.46) | 297.09 (64.84) | 313.29 (33.63) | 302.34 (59.65) | 312.76 (35.98) | 311.19 (45.11) | 297.61 (54.2) | 297.1 (43.59) | 302.16 (51.83) |
| R thalamus | 872.12 (110.88) | 846.66 (81.25) | 851.43 (83.95) | 851.52 (92.07) | 867.92 (105.39) | 847.07 (66.44) | 866.81 (86.71) | 838.07 (93.22) | 879.05 (83.72) | 831.57 (81.83) |
| R temporal pole | 45.84 (12.6) | 45.75 (12.02) | 45.69 (13.45) | 45.9 (10) | 51.82 (19.43) | 43.52 (12.83) | 48.53 (9.16) | 47.95 (13.67) | 39.6 (11.13) | 47 (13.03) |
| R transverse temporal | 75.45 (15.39) | 81.14 (15.04) | 79.77 (24.58) | 77.52 (15.03) | 76.5 (16.12) | 79.97 (15.96) | 80.43 (14.69) | 81.12 (19.69) | 78.96 (18.95) | 79.29 (15.94) |
| Global Efficiency | 2719.05 (106.9 9) | 2788.23 (96.9) | 2738.62 (118.0 6) | 2720.88 (146.8 1) | 2723.42 (90.21 ) | 2817.99 (99.72 ) | 2730.81 (111.1 1) | 2723.69 (127.8 4) | 2751.24 (97.1) | 2718.18 (113.1 2) |

#### Unharmonized data analyses

Table S7: Outcomes of network-based statistics analysis with unharmonized data

| HC > AN | Connection | T-statistic for between group difference | P-value for component[a] |
|---|---|---|---|
| | R thalamus - L hippocampus | 3.15 | 0.038 |
| | R thalamus - R hippocampus | 3.41 | |
| | R thalamus - R amygdala | 3.55 | |
| | L parahippocampal gyrus - R parahippocampal gyrus | 3.35 | |
| | R thalamus - R parahippocampal gyrus | 3.11 | |
| | R hippocampus - R postcentral gyrus | 3.26 | |
| | R paracentral gyrus - R postcentral gyrus | 3.34 | |
| | L cerebellum - R cerebellum | 3.11 | |
| | R parahippocampal gyrus - R cerebellum | 3.54 | |
| **AN > HC** | **Connection** | **T-statistic for between group difference** | **P-value for component** |
| | L lateral orbitofrontal gyrus - L pars opercularis | 3.29 | 0.018 |
| | L pars opercularis - L pars orbitalis | 3.58 | |
| | L caudal anterior cingulate – L pars triangularis | 3.12 | |
| | L pars orbitalis - L pars triangularis | 3.16 | |
| | L precentral gyrus – R putamen | 3.57 | |
| | L pars opercularis - R putamen | 3.22 | |
| | L precentral gyrus – L superior temporal gyrus | 3.31 | |
| | L pars opercularis - R lateral orbitofrontal gyrus | 3.28 | |
| | L lateral orbitofrontal gyrus - R rostral middle frontal gyrus | 3.98 | |
| | L lateral orbitofrontal gyrus - R rostral anterior cingulate cortex | 3.31 | |
| | L superior temporal gyrus - R putamen | 3.21 | |
| Models adjusted for age, motion, study, [a] P values Bonferonni-corrected for multiple comparisons, R: right, L:left | | | |

Table S8: Outcomes of between-group analyses of node strength calculated with unharmonized data

| | Strength | | | |
|---|---|---|---|---|
| **Node** | HC Mean (SD) | AN Mean (SD) | Effect of Diagnosis | P value |
| L nucleus accumbens | 18558.06 (4902.75) | 18167.33 (4685.57) | -457.35 [-996.49,81.8] | 1.00 |
| L amygdala | 28002.01 (6811.12) | 26609.86 (4800.53) | -702.6 [-1398.9,-6.31] | 1.00 |
| L bank of the superior temporal sulcus | 30908.79 (7295.44) | 32414.24 (7825.42) | 512.4 [-388.57,1413.36] | 1.00 |
| L caudal anterior cingulate | 28203.15 (5362.87) | 29193.31 (5476.48) | 591.4 [-69.12,1251.92] | 1.00 |
| L caudate | 99768.97 (16701.06) | 98790.67 (14758.58) | -627.99 [-2499.21,1243.23] | 1.00 |
| L Cerebellum | 50669.56 (12792.06) | 45578.9 (10830.9) | -2004.94 [-3316.51,-693.37] | 0.28 |
| L caudal middle frontal gyrus | 64931.64 (13160.15) | 65955.86 (9490.04) | 378.13 [-984.15,1740.41] | 1.00 |
| L cuneus | 42446.29 (8310.31) | 43323.61 (7941.82) | 427.27 [-562.08,1416.63] | 1.00 |
| L entorhinal | 22003.79 (5935.08) | 20561.67 (5816.47) | -570.34 [-1277.38,136.7] | 1.00 |
| L frontal pole | 8187.46 (3153.86) | 8172.39 (3600.43) | 172.32 [-212.35,557] | 1.00 |
| L fusiform | 63304.45 (10553.34) | 66514.15 (10331.44) | 1439.88 [216.8,2662.97] | 0.91 |
| L hippocampus | 80981.42 (10631.44) | 76136.86 (9403.45) | -2162.96 [-3378.72,-947.21] | 0.12 |
| L isthmus cingulate cortex | 39518.14 (5228.62) | 39713.02 (5370.17) | 270.03 [-364.51,904.57] | 1.00 |
| L insula | 103638.18 (13549.19) | 104301.78 (14712.42) | 171.28 [-1540.6,1883.16] | 1.00 |
| L inferior parietal lobule | 85545.93 (13645.42) | 88698.96 (12347.41) | 1337.3 [-192.08,2866.69] | 1.00 |
| L inferior temporal gyrus | 64807.88 (13191.55) | 69494.94 (12162.11) | 2206.3 [677.47,3735.14] | 0.37 |
| L lingual | 55029.18 (10215.03) | 54486.07 (8781.58) | -372.32 [-1451.68,707.04] | 1.00 |
| L lateral orbitofrontal | 65347.24 (9963.84) | 66294.54 (9375.84) | 648.77 [-524.27,1821.81] | 1.00 |
| L lateral occipital gyrus | 82407.66 (16505.63) | 86231.15 (13573.9) | 2110.9 [446.59,3775.22] | 0.80 |
| L medial orbitofrontal | 55110.33 (11289.52) | 54617.59 (10412.72) | -320.3 [-1629.27,988.68] | 1.00 |
| L middle temporal gyrus | 65243.23 (13253.12) | 66246.31 (12116.18) | 1056.6 [-298.14,2411.34] | 1.00 |
| L pallidum | 54286.58 (9905.73) | 52479.99 (10366.06) | -427.21 [-1592.81,738.39] | 1.00 |
| L paracentral | 45969.34 (8200.67) | 45949.12 (6096.01) | 20.91 [-830.18,872] | 1.00 |
| L parahippocampal | 34691.92 (6659.53) | 32025.75 (5348.64) | -1160.16 [-1879.28,-441.04] | 0.23 |
| L posterior cingulate cortex | 38588.19 (6656.18) | 39435.11 (5829.58) | 581.81 [-173.95,1337.57] | 1.00 |
| L precuneus | 79862.89 (13361.86) | 81161.49 (10709.95) | 565.5 [-770.12,1901.12] | 1.00 |
| L pericalcarine | 58077.4 (12957.45) | 58337.82 (13042.46) | 236.91 [-1296.15,1769.98] | 1.00 |
| L pars opercularis | 44534.35 (10458.24) | 46619.78 (9228.77) | 1227.15 [70.17,2384.12] | 0.98 |
| L pars orbitalis | 21792.06 (4531.07) | 23168.59 (4338.25) | 813.48 [276.11,1350.84] | 0.41 |
| L postcentral | 80374.29 (16096.74) | 81255.62 (10324.34) | 721.8 [-807.32,2250.91] | 1.00 |
| L precentral | 121487.47 (19446.12) | 123787.87 (10924.38) | 1346.86 [-522.99,3216.7] | 1.00 |
| L pars triangularis | 37689.02 (6965.84) | 38916.93 (6845.45) | 778.04 [-58.94,1615.03] | 1.00 |
| L putamen | 132502.28 (16407.71) | 135200.28 (17238.08) | 1280.41 [-803.06,3363.88] | 1.00 |
| L rostral anterior cingulate cortex | 33770.91 (6387.86) | 32389.53 (6051.56) | -499.62 [-1241.12,241.87] | 1.00 |
| L rostral middle frontal gyrus | 107928.87 (20835.95) | 111073.15 (18504.95) | 961.87 [-1328.74,3252.48] | 1.00 |
| L superior frontal gyrus | 149162.48 (23496.81) | 152823.18 (12476.87) | 2145.87 [-65.15,4356.9] | 1.00 |
| L supramarginal gyrus | 75378.51 (13172.1) | 75123.65 (9131.86) | 123.61 [-1220.87,1468.1] | 1.00 |
| L superior parietal lobule | 116685.05 (18257.83) | 121345.76 (15272.21) | 2145.74 [115.99,4175.49] | 0.99 |
| L superior temporal gyrus | 65479.37 (11773.62) | 66996.22 (9541.44) | 1062.91 [-120.37,2246.2] | 1.00 |
| L thalamus | 121895.45 (14050.93) | 119144.73 (13463.65) | -1139.65 [-2724.51,445.21] | 1.00 |
| L temporal pole | 13354.29 (4086.51) | 13371.52 (3998.44) | 148.27 [-336.18,632.73] | 1.00 |
| L transverse temporal | 15251.27 (2763.83) | 15591.65 (2898.93) | 170.44 [-160.03,500.91] | 1.00 |
| R nucleus accumbens | 21126.77 (5612) | 20859.52 (5117.06) | -232.17 [-859.81,395.47] | 1.00 |
| R amygdala | 32460.93 (7719.37) | 31410.13 (6909.62) | -507.72 [-1326.5,311.05] | 1.00 |

| | | | | |
|---|---|---|---|---|
| R bank of the superior temporal sulcus | 32292.71 (7870.74) | 33369.01 (8852.74) | 165.48 [-823.96,1154.93] | 1.00 |
| R caudal anterior cingulate | 31561.87 (6656.2) | 31755.1 (5888.15) | 337.19 [-408.93,1083.31] | 1.00 |
| R caudate | 106867.21 (17756.45) | 103698.89 (15577.06) | -1421.67 [-3350.92,507.58] | 1.00 |
| R Cerebellum | 50132.91 (13651.3) | 45229.68 (10697.54) | -1862.31 [-3214.21,-510.42] | 0.63 |
| R caudal middle frontal gyrus | 63493.36 (11891.57) | 64826.53 (10603.83) | 452.52 [-872.78,1777.83] | 1.00 |
| R cuneus | 45166.72 (7268.76) | 45279.48 (6777.15) | -2.32 [-860.92,856.28] | 1.00 |
| R entorhinal | 22577.82 (8396.87) | 20885.22 (6224.55) | -715.93 [-1575.84,143.98] | 1.00 |
| R frontal pole | 10049.2 (3763.04) | 10624.7 (4366.23) | 453.33 [-13.05,919.71] | 1.00 |
| R fusiform | 64547.11 (12595.42) | 68405.78 (12762.23) | 1308.09 [-150.57,2766.74] | 1.00 |
| R hippocampus | 87521.92 (13982.59) | 83535.73 (13871.67) | -1786.54 [-3415.99,-157.09] | 0.97 |
| R isthmus cingulate cortex | 36398.97 (6021.17) | 36711.2 (4646.28) | 308.34 [-325.39,942.08] | 1.00 |
| R insula | 107318.13 (21133.86) | 107971.72 (21365.97) | -338.88 [-2738.53,2060.78] | 1.00 |
| R inferior parietal lobule | 92767.08 (13074.92) | 95632.61 (11917.31) | 746.14 [-685.48,2177.76] | 1.00 |
| R inferior temporal gyrus | 65619.34 (13238.26) | 67872.46 (11475.76) | 1127.65 [-368.31,2623.62] | 1.00 |
| R lingual | 59280.27 (9999.32) | 58676.46 (8060.15) | -396.02 [-1438.06,646.01] | 1.00 |
| R lateral orbitofrontal | 71429.63 (11905.63) | 71945.88 (11940.98) | 242.92 [-1172.6,1658.45] | 1.00 |
| R lateral occipital gyrus | 84694.92 (11611.34) | 89562.48 (11259.95) | 2183.24 [822.16,3544.32] | 0.26 |
| R medial orbitofrontal | 48590.33 (8737.83) | 48514.76 (9062.35) | -6.9 [-1110.06,1096.25] | 1.00 |
| R middle temporal gyrus | 70536.12 (13172.71) | 70836.81 (11962.1) | 607.05 [-780.03,1994.13] | 1.00 |
| R pallidum | 55197.78 (11523.13) | 55034.93 (11344.51) | 266.42 [-1023.34,1556.19] | 1.00 |
| R paracentral | 50347.97 (8136.49) | 49893.63 (5472.24) | -242.31 [-1082.14,597.52] | 1.00 |
| R parahippocampal | 35501.49 (7131.1) | 33554.78 (6479.03) | -915.19 [-1726.68,-103.69] | 0.95 |
| R posterior cingulate cortex | 39298.95 (6388.33) | 39480.13 (6170.34) | 308.16 [-443.24,1059.56] | 1.00 |
| R precuneus | 81585.94 (11535.97) | 82841.12 (10213.28) | 184.28 [-1056.3,1424.85] | 1.00 |
| R pericalcarine | 61792.85 (11086.04) | 61926.56 (11591.96) | -87.73 [-1467.27,1291.81] | 1.00 |
| R pars opercularis | 43485.29 (8830.31) | 44069.51 (9338.78) | 347.25 [-645.2,1339.7] | 1.00 |
| R pars orbitalis | 25910.56 (5302.11) | 26519.79 (5050.35) | 367.56 [-255.55,990.67] | 1.00 |
| R postcentral | 82008.71 (14285.42) | 81374.24 (11993.52) | -123.05 [-1555.71,1309.62] | 1.00 |
| R precentral | 122339.42 (16923.12) | 123066.56 (11950.01) | 530.94 [-1149.75,2211.62] | 1.00 |
| R pars triangularis | 40732.84 (7013.96) | 41491 (8248.24) | 521.41 [-404.74,1447.55] | 1.00 |
| R putamen | 136515.13 (17522.39) | 141782.09 (23783.08) | 2255.16 [-194.6,4704.92] | 1.00 |
| R rostral anterior cingulate cortex | 27748.81 (5748.13) | 27464.41 (5370.59) | 8.89 [-632.09,649.87] | 1.00 |
| R rostral middle frontal gyrus | 113512.5 (22645.11) | 116312.28 (22411.16) | 757.95 [-1758.83,3274.74] | 1.00 |
| R superior frontal gyrus | 144946.08 (20391.87) | 148268.73 (13761.88) | 2001.01 [-53.37,4055.39] | 1.00 |
| R supramarginal gyrus | 72279.22 (11672.04) | 72680.45 (10625.43) | 205.48 [-1090.84,1501.81] | 1.00 |
| R superior parietal lobule | 114245.61 (17140.24) | 118873.86 (15029.28) | 1582.71 [-296.85,3462.28] | 1.00 |
| R superior temporal gyrus | 61210.75 (9643.29) | 62030.73 (11678.87) | 749.74 [-500.87,2000.34] | 1.00 |
| R thalamus | 113774.93 (13087.41) | 109231.49 (11710.13) | -2014.32 [-3429.77,-598.87] | 0.54 |
| R temporal pole | 14886.57 (6100.13) | 14687.5 (4677.97) | 4.41 [-619.56,628.39] | 1.00 |
| R transverse temporal | 13735.7 (2514.06) | 13492.99 (2220.77) | -107.84 [-395.04,179.36] | 1.00 |

All models adjusted for age, motion, study. [a] P values permutation-corrected for multiple comparisons, R: right, L:left

Table S9: Outcomes of between-group analyses of node efficiency calculated with unharmonized data

| | Efficiency | | | |
|---|---|---|---|---|
| **Node** | HC Mean (SD) | AN Mean (SD) | Effect of Diagnosis | P value[a] |
| L nucleus accumbens | 202.29 (36.78) | 196.72 (35.83) | -3.8 [-8.11,0.52] | 1.00 |
| L amygdala | 270.37 (56.07) | 267.43 (43.81) | -1.63 [-7.49,4.23] | 1.00 |
| L bank of the superior temporal sulcus | 246.61 (70.64) | 248.71 (38.26) | -0.06 [-6.83,6.71] | 1.00 |
| L caudal anterior cingulate | 299.6 (49.47) | 307.36 (49.42) | 4.63 [-1.31,10.57] | 1.00 |
| L caudate | 642.05 (74.16) | 643.88 (81) | 0.75 [-8.76,10.25] | 1.00 |
| L Cerebellum | 251.47 (51.18) | 238.25 (46.71) | -5.94 [-11.78,-0.09] | 1.00 |
| L caudal middle frontal gyrus | 441.6 (73.72) | 446.22 (68.22) | 3.46 [-5.2,12.12] | 1.00 |
| L cuneus | 274.5 (54.78) | 283.5 (56.42) | 3.45 [-2.91,9.8] | 1.00 |
| L entorhinal | 285.25 (84.34) | 283.39 (74.38) | -1.98 [-11.32,7.35] | 1.00 |
| L frontal pole | 177.66 (52.36) | 176.56 (57.22) | 0.11 [-6.65,6.88] | 1.00 |
| L fusiform | 394.47 (110.09) | 394.06 (66.63) | -0.94 [-11.81,9.92] | 1.00 |
| L hippocampus | 468.69 (63.95) | 441.92 (57.45) | -11.12 [-18.35,-3.88] | 0.30 |
| L isthmus cingulate cortex | 381.54 (49.27) | 380.59 (49.49) | 0.97 [-4.73,6.67] | 1.00 |
| L insula | 691.7 (71.81) | 697.42 (74.85) | 2.23 [-6.64,11.1] | 1.00 |
| L inferior parietal lobule | 527.98 (91.45) | 540.13 (59.51) | 6.24 [-2.86,15.34] | 1.00 |
| L inferior temporal gyrus | 407.41 (84.58) | 415.65 (55.35) | 3.69 [-4.92,12.3] | 1.00 |
| L lingual | 359.87 (75.09) | 365.81 (57.51) | 1.8 [-6.11,9.7] | 1.00 |
| L lateral orbitofrontal | 438.59 (62.72) | 448.38 (58.75) | 5.84 [-1.4,13.08] | 1.00 |
| L lateral occipital gyrus | 449.84 (83.24) | 451.41 (55.37) | 1.76 [-6.44,9.96] | 1.00 |
| L medial orbitofrontal | 389.36 (62.08) | 387.18 (61.65) | -0.66 [-8.02,6.71] | 1.00 |
| L middle temporal gyrus | 376.61 (87.28) | 372 (53.02) | 0.42 [-7.89,8.73] | 1.00 |
| L pallidum | 470.55 (62) | 459.73 (66.07) | -1.87 [-9.05,5.31] | 1.00 |
| L paracentral | 344.27 (52.82) | 351.86 (49.48) | 4.53 [-1.66,10.71] | 1.00 |
| L parahippocampal | 252.77 (49.47) | 238.19 (37.92) | -6.88 [-12.04,-1.72] | 0.65 |
| L posterior cingulate cortex | 362.58 (64.35) | 371.93 (62.09) | 5.9 [-1.81,13.6] | 1.00 |
| L precuneus | 586.63 (70.77) | 581.29 (58.95) | -1.18 [-8.3,5.95] | 1.00 |
| L pericalcarine | 300.59 (59.44) | 317.86 (65.8) | 8.48 [1.14,15.81] | 0.90 |
| L pars opercularis | 393.64 (69.47) | 409.03 (54.76) | 9.33 [1.86,16.8] | 0.76 |
| L pars orbitalis | 221.22 (45.25) | 230.32 (34.48) | 4.82 [0.1,9.53] | 0.99 |
| L postcentral | 448.27 (49.99) | 456.37 (53.72) | 6.19 [0,12.37] | 1.00 |
| L precentral | 657.02 (104.85) | 674.42 (71.76) | 11 [0.3,21.71] | 0.99 |
| L pars triangularis | 324.01 (44.85) | 333.46 (40.22) | 5.66 [0.48,10.83] | 0.98 |
| L putamen | 811.95 (77.57) | 827.15 (81.4) | 9.6 [0.04,19.17] | 0.99 |
| L rostral anterior cingulate cortex | 340.71 (60.36) | 334.22 (59.98) | -2.35 [-9.12,4.41] | 1.00 |
| L rostral middle frontal gyrus | 641.11 (91.05) | 647.18 (62.15) | 3.5 [-5.53,12.54] | 1.00 |
| L superior frontal gyrus | 865.71 (110.36) | 872.43 (77.68) | 7.24 [-3.55,18.03] | 1.00 |
| L supramarginal gyrus | 476.42 (64.58) | 480.59 (58.14) | 3.56 [-3.92,11.05] | 1.00 |
| L superior parietal lobule | 696.87 (89.7) | 715.22 (78.66) | 11.21 [2.05,20.37] | 0.85 |
| L superior temporal gyrus | 429.86 (68.53) | 428.37 (46.93) | 0.28 [-6.64,7.21] | 1.00 |
| L thalamus | 943.97 (85.18) | 928.68 (87.36) | -5.81 [-16.06,4.45] | 1.00 |
| L temporal pole | 299.45 (162.24) | 287.87 (101.96) | -5.98 [-21.92,9.95] | 1.00 |
| L transverse temporal | 180.32 (46.03) | 185.14 (28.43) | 1.96 [-2.62,6.53] | 1.00 |
| R nucleus accumbens | 219.75 (52.01) | 220.51 (37.29) | -0.45 [-5.87,4.96] | 1.00 |
| R amygdala | 291.37 (60.4) | 289.89 (50.19) | -1.07 [-7.33,5.19] | 1.00 |
| R bank of the superior temporal sulcus | 248.39 (73.75) | 251.08 (51.41) | -0.71 [-7.8,6.37] | 1.00 |

| | | | | |
|---|---|---|---|---|
| R caudal anterior cingulate | 322.64 (59.99) | 322.14 (44.76) | 2.09 [-4.09,8.26] | 1.00 |
| R caudate | 695.92 (110.79) | 670.8 (86.4) | -10.77 [-22.36,0.83] | 1.00 |
| R Cerebellum | 247.8 (54.32) | 236.64 (42.69) | -5.29 [-10.97,0.39] | 1.00 |
| R caudal middle frontal gyrus | 432.26 (76.89) | 438.04 (58.25) | 2.97 [-4.82,10.75] | 1.00 |
| R cuneus | 279.2 (46.81) | 289.06 (45.35) | 3.63 [-1.76,9.02] | 1.00 |
| R entorhinal | 298.14 (112.21) | 303.65 (105.61) | 2.99 [-9.12,15.11] | 1.00 |
| R frontal pole | 191.54 (49.13) | 189.24 (46.77) | -0.62 [-5.75,4.51] | 1.00 |
| R fusiform | 400.78 (98.43) | 410.81 (74.91) | 2.73 [-7.46,12.92] | 1.00 |
| R hippocampus | 526.57 (87.53) | 505.52 (72.26) | -8 [-17.28,1.28] | 1.00 |
| R isthmus cingulate cortex | 343.5 (55.96) | 344.37 (41.05) | 1.84 [-3.91,7.6] | 1.00 |
| R insula | 720.88 (154.13) | 702.66 (99.6) | -11.51 [-26.32,3.3] | 1.00 |
| R inferior parietal lobule | 570.56 (82.39) | 578.1 (62.3) | 4 [-4.84,12.85] | 1.00 |
| R inferior temporal gyrus | 400.33 (97.9) | 395.63 (53.64) | -1.25 [-10.25,7.75] | 1.00 |
| R lingual | 381.23 (66.03) | 373.89 (54.22) | -5.03 [-12.14,2.08] | 1.00 |
| R lateral orbitofrontal | 464.75 (90.7) | 467.8 (64.33) | 2.89 [-5.98,11.76] | 1.00 |
| R lateral occipital gyrus | 461.24 (92.4) | 463.94 (50.51) | 2.08 [-6.34,10.5] | 1.00 |
| R medial orbitofrontal | 378.19 (78.23) | 365.77 (76.02) | -5.12 [-13.52,3.29] | 1.00 |
| R middle temporal gyrus | 404.83 (112.41) | 391.83 (53.13) | -3.2 [-13,6.59] | 1.00 |
| R pallidum | 469.94 (90.63) | 467.97 (64.83) | 1.6 [-7.54,10.74] | 1.00 |
| R paracentral | 373.73 (62.55) | 373.54 (53.7) | 1.07 [-5.85,7.99] | 1.00 |
| R parahippocampal | 263.1 (58.74) | 252.86 (45.18) | -4.8 [-10.78,1.17] | 1.00 |
| R posterior cingulate cortex | 371.5 (66.25) | 375.52 (60.7) | 3.75 [-3.96,11.46] | 1.00 |
| R precuneus | 584.17 (85.35) | 584.59 (53.68) | -0.08 [-8.24,8.09] | 1.00 |
| R pericalcarine | 306.33 (59.64) | 318.47 (53.59) | 4.33 [-2.21,10.86] | 1.00 |
| R pars opercularis | 374.17 (67.87) | 374.82 (51.2) | 1.02 [-5.76,7.79] | 1.00 |
| R pars orbitalis | 242.78 (52.44) | 246.56 (44.85) | 2.83 [-2.56,8.22] | 1.00 |
| R postcentral | 449.37 (75.63) | 445.06 (50.73) | -0.57 [-8,6.86] | 1.00 |
| R precentral | 653.75 (103.55) | 661.45 (63.23) | 6.6 [-3.41,16.61] | 1.00 |
| R pars triangularis | 344.97 (52.89) | 341.17 (52.4) | -0.3 [-6.41,5.82] | 1.00 |
| R putamen | 837.75 (131.85) | 854.15 (89.36) | 10.32 [-3.2,23.84] | 1.00 |
| R rostral anterior cingulate cortex | 331.2 (59.69) | 327.12 (56.76) | -0.27 [-6.63,6.09] | 1.00 |
| R rostral middle frontal gyrus | 657.76 (96.13) | 668.75 (80.22) | 5.98 [-4.1,16.06] | 1.00 |
| R superior frontal gyrus | 845.21 (111.76) | 854.15 (83.41) | 8.65 [-2.92,20.22] | 1.00 |
| R supramarginal gyrus | 471.68 (71.63) | 463.53 (58.53) | -2.65 [-10.24,4.93] | 1.00 |
| R superior parietal lobule | 698.76 (112.36) | 708.54 (67.44) | 5.61 [-5.34,16.55] | 1.00 |
| R superior temporal gyrus | 419.59 (100.8) | 414.51 (62.09) | -1.41 [-10.7,7.89] | 1.00 |
| R thalamus | 901.04 (115.38) | 867.34 (83.56) | -14.88 [-26.86,-2.91] | 0.80 |
| R temporal pole | 360.46 (228.93) | 353.17 (175.41) | -6.07 [-28.64,16.51] | 1.00 |
| R transverse temporal | 177.33 (39.19) | 175.21 (32.85) | -1.57 [-5.65,2.5] | 1.00 |

All models adjusted for age, motion, study. [a] P values permutation-corrected for multiple comparisons, R: right, L:left

**Subtype analyses**

Table S10: Differences in structural connectivity metrics between AN subtypes

| Structural connectivity metric | **B [95 %CI]** | P-value |
|---|---|---|
| Subcortical component connectivity (mean streamline count) | 249.03 [-19.12, 517.18] | 0.0 |
| Cortical component connectivity (mean streamline count) | 0.82 [-21.29,22.92] | 0.94 |
| Strength of left hippocampus | 532.37 [-2630.00, 3694.75] | 0.74 |

Models adjusted for age, motion, study, body mass index (BMI). The BMI variable combined scaled BMI scores calculated in adults of the sample (aged over 19 years) with scaled BMI percentile scores calculated in adolescents (aged 19 years and below).

**Subsample analyses**

Table S11: Results of analyses testing between-group differences in structural connectivity metrics in different age-groups

| Structural connectivity metric | Participants aged 19 and under (N = 60 AN, 65 HC) | | Participants aged 20 and over (N = 87 AN, 54 HC) | |
|---|---|---|---|---|
| | B [95 %CI] | P-value | B [95 %CI] | P-value |
| Subcortical component connectivity (mean streamline count) | -307.78 [-451.64,-163.92] | <0.001 | -445.09 [-620.60,-269.57] | <0.001 |
| Cortical component connectivity (mean streamline count) | 26.12 [15.16, 37.09] | <0.001 | 15.49 [4.45,26.52] | <0.001 |
| Strength of left hippocampus | -3077.26 [-4665.35,-1489.17] | <0.001 | -2062.91 [-3942.71,-183.10] | 0.03 |

Models adjusted for age, motion, and study.
